# Cultural transmission, competition for prey, and the evolution of cooperative hunting

**DOI:** 10.1101/2022.12.14.520505

**Authors:** Talia Borofsky, Marcus W. Feldman, Yoav Ram

## Abstract

Although cooperative hunting (CH) is widespread among animals, its benefits are unclear. When rare, CH may allow predators to escape competition and access “big prey” (BP). However, a lone CH predator cannot such catch food. Cultural transmission may allow CH to spread fast enough that cooperators can find hunting partners, but competition for BP may increase. We construct a one-predator, two-prey model in which the predators either learn to hunt “small prey” (SP) alone, or learn to hunt BP cooperatively. The predators first learn vertically and then choose partners from which they learn horizontally with probability *H*. CH predators only catch the BP if their partner is cooperative. We find that without horizontal learning, CH cannot evolve when initially rare. Together, a high probability of horizontal learning and competition for the SP allow CH to evolve. However, CH can only fix in the predator population if the BP is very abundant. Furthermore, a mutant that increases horizontal learning can invade whenever CH is present but not fixed, because horizontal learning allows predators to match their strategies, avoiding the situation in which a cooperator cannot find a partner. While competition for prey is important for determining the degree of CH that evolves, it is not enough for CH to emerge and spread; horizontal cultural transmission is essential. Future models may explore factors that control how horizontal transmission influences cooperative predation, and vice versa. Lessons from our model may be useful in conservation efforts and wildlife reintroduction programs.

## 1 Introduction

Competition for resources is a potent ecological force that could influence the evolution of cooperative traits (Frank, 2019). However, the effect of competition on the evolution of altruism depends on the type of altruism in question. Van Dyken and Wade (2012a; 2012b) group the fitness components that can be donated or shared by the altruist into three categories: resources, survival, and fecundity. Although previous research has suggested that local competition selects against altruism (Frank, 2019), the story changes if we differentiate between types of altruism based on the relevant fitness components; while local competition selects against fecundity altruism, such as cooperative breeding, it selects for resource altruism, such as cooperative hunting (Van Dyken and Wade, 2012a). In fact, resource altruism may decrease local competition and in the process drive the evolution of survival or fecundity altruism (Van Dyken and Wade, 2012b). Cooperative hunting, also called group hunting, is a type of resource altruism in which multiple predators hunt prey together and share what they catch. It is performed by a wide variety of animals such as spiders, lions, wolves, whales, dolphins, chimpanzees, and of course, humans (Alvard et al., 2002; Boesch, 1994; Mangel et al., 1988; Uetz, 1992; Whitehead and Rendell, 2014). Group-hunting behaviors can be quite complex, and for many species the extent to which predators must learn them is unclear. For example, type B killer whales (*Orcinus orca*) have been documented to hunt seals on the Antarctic ice pack by “wave washing”: multiple orcas simultaneously lunge toward an ice flow and produce a wave that pushes the seal into the water (Pitman and Durban, 2012). It may be difficult for an individual orca to discover this hunting behavior because waves generated by one or two orcas rarely succeed at pushing seals into the water (Pitman and Durban, 2012). However, it is plausible that the wave washing behavior first emerged when one orca attempted to lunge at a seal on an ice flow while another orca watched and then joined in. By acting together, they may have created a wave large enough to wash a seal into the water.Although scientists have not been able to directly observe social learning of cooperative hunting by killer whales, it might be inferred because (1) mothers and their offspring exhibit similar behaviors (Rendell and Whitehead, 2001), (2) there are large foraging behavior differences between pods, but little variation within pods (Rendell and Whitehead, 2001), (3) there are several known instances of social learning of foraging skills in Cetaceans (Allen et al., 2013; Krützen et al., 2005; Mann et al., 2012; Whitehead, 2017), including strong evidence for horizontal and oblique learning of a foraging behavior in dolphins (Wild et al., 2020) and humpback whales (Weinrich et al., 1992), and (4) in captivity, killer whales have demonstrated the ability to imitate actions performed by other killer whales (Abramson et al., 2013). In fact, it has been suggested that the cultural transmission of complex cooperative foraging behaviors promoted sympatric speciation between killer whale ecotypes (Foote et al., 2016; Riesch et al., 2012; Whitehead, 2017).

Cooperative hunting is not restricted to group-living animals. The Malagassy fossa (*Cryptoprocta ferox*) is one of the few known animals that are generally solitary but sometimes hunt in groups. This mammalian predator lives in Madagascar, and groups of 2 - 3 male fossas have been observed to cooperatively hunt a type of large lemur called Verraux’s sifaka (*Propithecus verrauxi*). To catch a sifaka, each fossa takes turns chasing it up a tree until the lemur has been chased into short trees where it can be caught (Lührs and Dammhahn, 2010). Fossas that hunt in groups tend to be larger and have more success in mating with females (Lührs et al., 2013). We can imagine a simple origin for this complex behavior: one fossa was socially attentive enough to observe another attempting to chase down a lemur, joined in the hunt, and then benefited from the prey being large enough to share. In fact, a species of giant lemur lived in Madagascar until recently and would have made sharing especially easy; this extinct lemur may have facilitated the spread of cooperative hunting in fossas (Lührs and Dammhahn, 2010).

Here, we aim to understand what behavioral and ecological characteristics allow cooperative hunting to emerge. As in the examples of the fossas and even the killer whales, cooperative hunting may have arisen simply from (1) a motivation to hunt a certain prey together, and (2) social attentiveness. Thus, cooperative hunting may not require a complicated communication system. As a proof of concept, it has been shown that seemingly coordinated group hunting behavior by wolves can be recreated in agent-based simulations with agents following very simple rules: move towards the prey, keep a safe distance from the prey, and do not run into other wolves (Muro et al., 2011).

The benefit of hunting in groups is not always clear. Previous models suggested that cooperative hunting can only be an evolutionarily stable strategy (ESS) against solitary hunting (Packer and Ruttan, 1988) if cooperative predators have a much higher payoff (i.e., from a higher rate of prey encounter, higher hunting success rate, or lower energetic cost) than solitary predators. This implies that if a predator lives in an environment with strong competition for existing resources, but with an abundance of a prey type that is costly to capture individually—for instance, because the prey can jump between trees, as in the sifaka discussed earlier— then the predator population may benefit from cooperative hunting.

Furthermore, a cooperative hunting behavior may spread more easily in a population if it is transmitted via social learning. A cooperative hunter cannot catch prey if it only interacts with solitary hunters, but if predators learn from their interaction partners, they can match behaviors so either both cooperate together or both hunt separately. In a model by Cohen et al. (2021), increased cooperation evolved when individuals learned from their interaction partners. Moreover, in cultural evolutionary models, behaviors that are detrimental to fitness may still fix in a population if they are culturally transmitted (i.e. transmitted via social learning) reliably enough (Cavalli-Sforza and Feldman, 1981). For example, cooperative hunting may at first be beneficial if a “small prey” (SP) that is hunted solitarily is depleted, while a “big prey” (BP) that is hunted cooperatively is abundant. However, cultural transmission could spread cooperative hunting fast enough that it continues to be present in the population even when the BP are depleted.

We investigate how the population dynamics of prey and social transmission of hunting strategies can influence the evolution of cooperative hunting. In comparison to cooperative predator defense by prey, few predator-prey models with dynamic population sizes have been developed for cooperative hunting by predators (but see Alves and Hilker, 2017; Banerjee et al., 2020; Berec, 2010; Van Dyken and Wade, 2012b). None of these models have examined the interaction between the evolution of the cooperative trait and social learning. Here, we develop a model in which predators can learn a hunting behavior that targets either the small prey, which is hunted solitarily, or the big prey, which must be hunted cooperatively. In our model, being a cooperator can be risky: the benefits are high (the BP has a larger effect on fitness than the SP, even after being shared), but the predators may not find a cooperative hunting partner to hunt the big prey. We assume predators first learn vertically, i.e., from their parents, and then may learn horizontally, i.e., from peers of the same generation, as in Cohen et al. (2021). Our hypotheses are:

(H1) Horizontal social learning selects for cooperative hunting.

(H2) Strong competition for small prey selects for cooperative hunting.

(H3) Depletion of the big prey limits the evolution of cooperative hunting.

(H4) If the previous hypotheses are true, then environmental conditions that select for cooperative hunting (i.e. there is more competition for small prey than big prey and the big prey is more beneficial) will also select for horizontal learning.

## 2 Model

Consider a population of predators in which each predator can hunt one of two types of prey: small prey (SP), which can be caught by solitary hunting (SH), and big prey (BP), which can be caught by cooperative hunting (CH). Both prey types are assumed to have the same carrying capacity, and their normalized densities are *r*_*s*_, and *r*_*b*_, respectively (the conversion from population densities to normalized densities is in Appendix A1).

### Predatordynamics

Assume, for simplicity, that predators that learn to hunt solitarily only hunt the SP and those that learn to hunt cooperatively only hunt the BP. The frequencies of behaviors *CH* and *SH* in the predator population and *p* are *q* = 1 − *p*, respectively. The densities of the big prey and small prey, normalized to be between 0 and 1, are *r*_*b*_ and *r*_*s*_, respectively. A cooperative predator successfully catches the BP if another cooperative predator is present.

Predators have the following lifecycle: (1) Offspring learn a hunting behavior by vertical cultural transmission from their parents. (2) Adults learn horizontally (i.e., from members of their own generation) with probability *H*, where 0 ≤ *H* ≤ 1. At every learning interaction between a pair of predators, each of the predators has a 50% chance to be the demonstrator while the other is the learner. The process of horizontally learning to be cooperative may only require the learner to be socially attentive to conspecifics hunting, and tolerant enough of conspecifics to be willing to join in. Likewise, horizontal learning of a solitary behavior may be quite simple; the cooperative hunter sees that a conspecific is not available to cooperate and is hunting SP, and from observation recognizes (or learns) that the SP is food.

During horizontal learning, cooperative predators that encounter other cooperative predators continue to be cooperative. The frequency of encounters between cooperative and solitary predators is 2*p*(1 − *p*), the probability of horizontal learning is *H*, and since there is an unbiased coinflip determining which individual is the demonstrator, the probability that a solitary hunter learns to be cooperative is *Hp*(1 − *p*). Hence the frequency 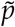 of CH among adults in the current generation (after vertical and horizontal transmission) is

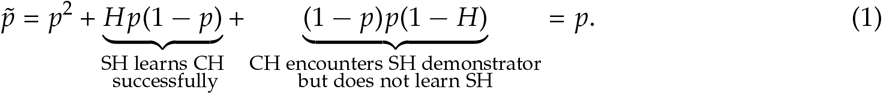

Therefore, the probability of horizontal learning *H* does not affect the actual frequency of cooperative hunting because the frequency at which cooperative individuals learn horizontally to be solitary is the same as the frequency of solitary individuals that learn horizontally to be cooperative.

Adults grow and become parents at a rate proportional to the amount of food they ate. If two cooperative hunters interact, the fitness of each is 1 + *r*_*b*_*b*_*c*_/2, where *b*_*c*_ is the energetic benefit of the BP, and the factor of 1/2 is due to food sharing. If a CH predator interacts with an SH predator, it does not catch prey, and its fitness is 1. A solitary hunter always has fitness 1 + *r*_*s*_*b*_*s*_, where *b*_*s*_ is the energetic benefit of the SP. We assume that if BP and SP have the same normalized densities, i.e. *r*_*s*_ = *r*_*b*_, then the BP gives a higher energetic benefit even after sharing compared to the SP alone, i.e., *b*_*c*_/2 > *b*_*s*_. We assume that catching BP is more beneficial than SP so that cooperating is beneficial when CH is common and BP is abundant. This assumption allows the model to focus on which ecological conditions (i.e. prey population dynamics) and cultural characteristics (i.e. frequency of CH, probability of horizontal transmision) limit or enable the evolution of CH. The payoff matrix for the interactions and a diagram of the model is in Fig. 1.

**Figure 1:**
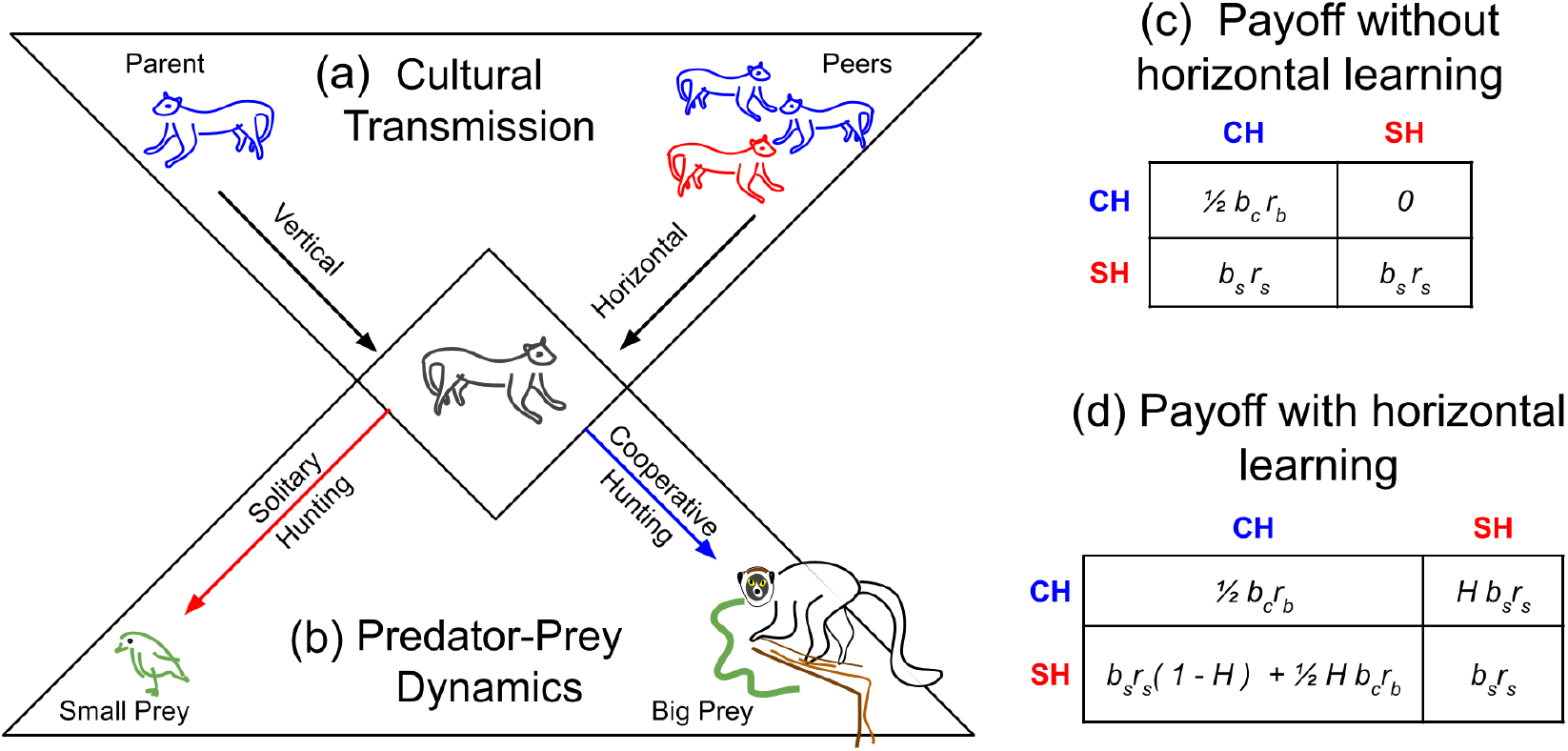
An illustration of the model structure. **(a)** The cultural transmission, or social learning dynamics. Individuals learn vertically from their parent (left) and then learn horizontally from other peers (right). **(b)** The predator-prey dynamics of the model. After learning, the predators can hunt cooperatively for the big prey (right) or hunt solitarily for the small prey (bottom). **(c)** The payoff matrix without horizontal learning, *H* = 0. **(d)** The payoff matrix with horizontal learning, *H* > 0.

Unlike Cohen et al. (2021), we assume that the predators interact after horizontal transmission. Thus, although we have shown that horizontal learning does not change the net frequency of cooperative hunting in the population (because learning the solitary and cooperative behaviors is symmetric), it rearranges cooperators and solitary hunters so that the the probability that a cooperator can find a partner with which to hunt the big prey increases. The frequencies *p*′ and q′ = 1 − *p*′ of parents in the current generation are given by the following recursions:

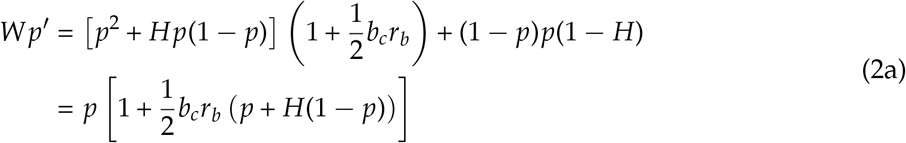

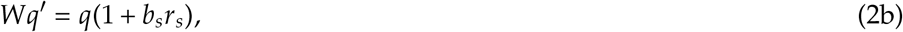

where *W* is the mean fitness of the predator population, i.e., the sum of the right sides of Eqs. (2a) and (2b), namely

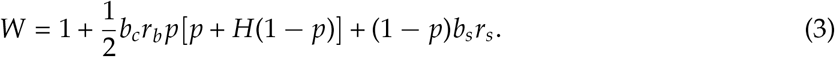

### Prey dynamics

We also track the prey population dynamics. The population dynamics in terms of normalized densities 0 ≤ *r*_*s*_, *r*_*b*_ ≤ 1 are derived in Appendix A1. The recursions are

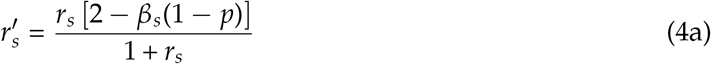

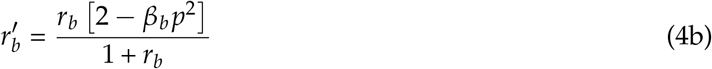

where *β*_*s*_, *β*_*b*_ are the depletion constants of the SP and BP, respectively. Assume random encounters between predators and prey, and that the encounter rate with a pair of predators is smaller than the encounter rate with one predator, i.e. *β*_*b*_ < *β*_*s*_ < 1.

## 3 Results

Note that prey equilibria are 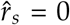 or 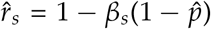 and 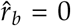 or 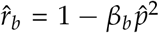, where 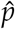 is the equilibrium frequency of cooperators in the predator population.

### Result 3.1.

*Neither prey can go extinct, i*.*e*. 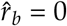 or 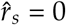 *are not locally stable*.

*Proof*. From (4a), if *r*_*s*_ is very small, 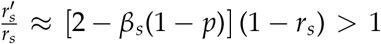 and from (4b), if *r*_*b*_ is very small, then also 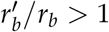.

### 3.1. Evolution of Cooperative Hunting without Horizontal Learning (H = 0)

Without horizontal transmission, *H* = 0, the change in frequency of cooperative hunting is derived by setting *H* = 0 in Eqs. 2. The equilibrium frequency of CH is found by setting 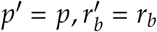 and 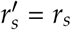 in Eqs. 2. The equilibria 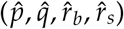 are the following:

1. Hunting strategies are polymorphic in the population and both prey types are present, i.e.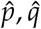 are found by dividing (2*a*) by (2*b*), 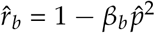, and 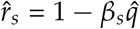.
2. SH fixes and both prey are present, i.e. 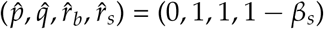.
3. CH fixes and both prey are present, i.e. 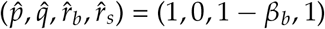
4. SP or BP go extinct (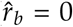or 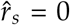: we do not discuss these further because they are not locally stable (Result 3.1)

Below, we analyze local stability and existence conditions for these equilibria.

#### Result 3.2.

*Suppose both types of prey are present at equilibrium densities* 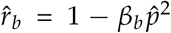 *and* 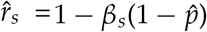. *Then there is one polymorphic predator equilibrium* 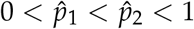 *if β*_*b*_ < *γ*_1_, *where*

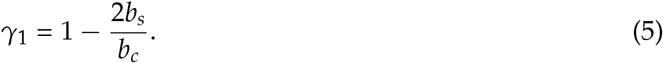

*If β*_*b*_ ≥ *γ*_1_, *there are two polymorphic predator equilibria*, 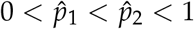, *if γ*_2_ < *β*_*b*_ < *γ*_3_, *where*

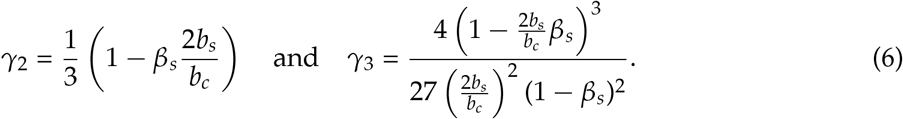

### Otherwise there is no polymorphic equilibrium

The proof is in Appendix A2.1.

#### Result 3.3.

*CH cannot increase when rare if* 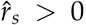, *i*.*e. the equilibrium* 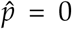, *is locally stable for* 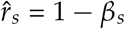 *and either* 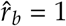 *or* 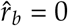.

*Proof*. Near 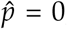, from (2a), 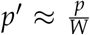 (neglecting higher order terms in *p*), where with *p* small, *W* ≈ 1 + *b*_*s*_*r*_*s*_. Thus *p*′ < *p* and 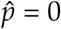 is locally stable.

#### Result 3.4.

*If both types of prey are present*, 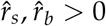 *and β*_*b*_ < *γ*_1_, *then CH can fix if it starts at a high enough frequency, i*.*e. the equilibrium* 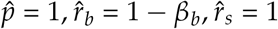 *is locally stable*.

*Proof*. Near the equilibrium 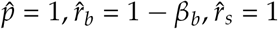, the Jacobian *J*\* for its local stability is given by Equations (A2.6) and (A2.7) in Appendix A2. The three eigenvalues of *J*\* are 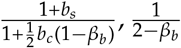, and 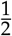. The first eigenvalue is less than one if 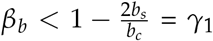 and the remaining eigenvalues are always less than one because *β*_*b*_ < 1.

Trajectories of the cooperative hunting frequency over time with parameter combinations for which there are no polymorphic equilibria, one polymorphic equilibrium, and two polymorphic equilibria are shown in Figs 2a - 2c and parameter regions for which different equilibria exist are shown in Appendix Figures A5.1 and A5.2.

**Figure 2:**
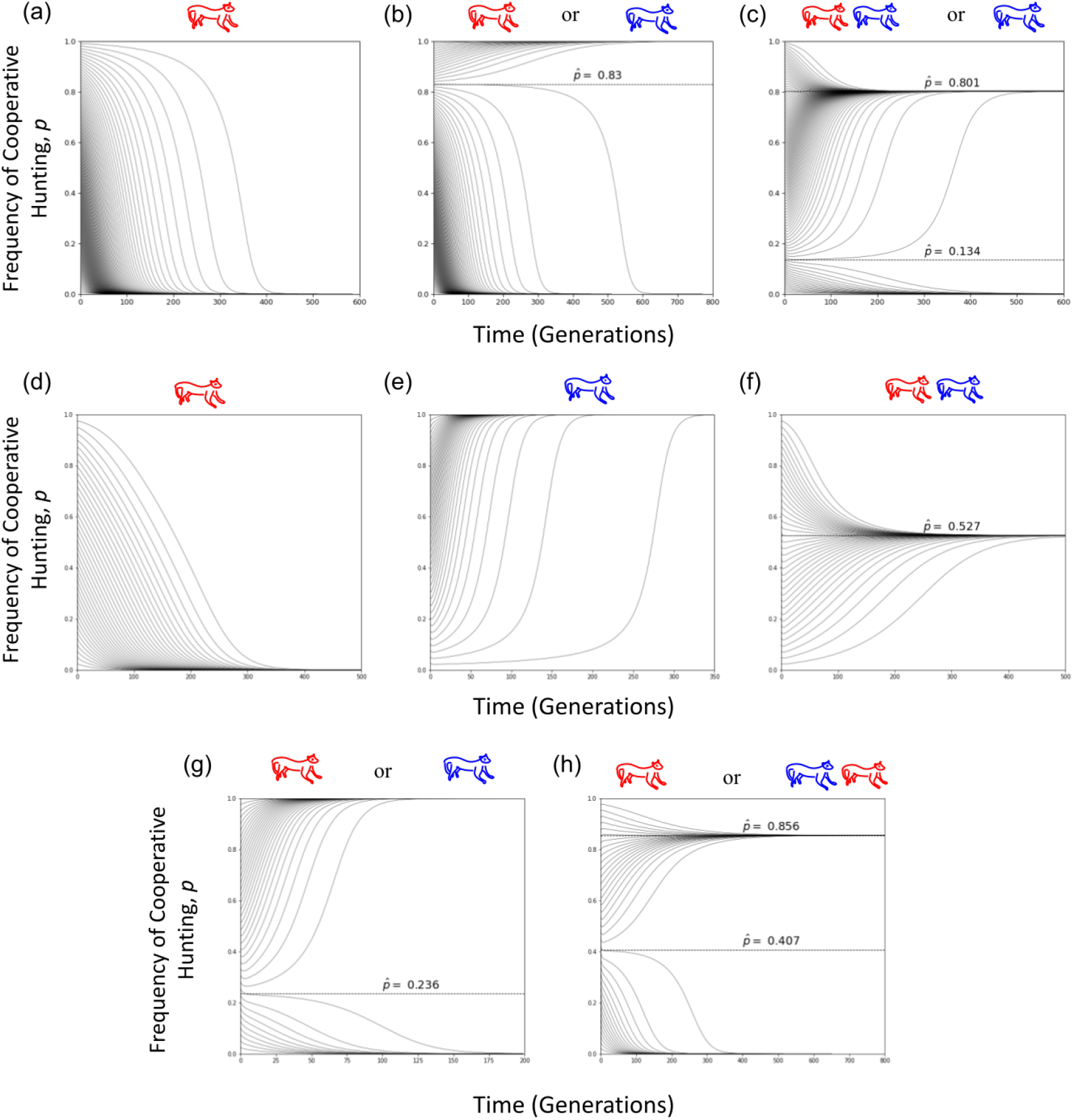
Trajectories of the frequency of cooperative hunting (CH), *p*, (y-axis) over time (generations) under vertical transmission only, *H* = 0 (panels (a) - (c)) or with horizontal transmission *H* > 0 (panels (d) - (h)). Dotted lines indicate polymorphic equilibria, i.e. both CH and SH are present. The initial prey values are 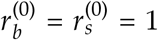, but different initial *p* values are chosen in the interval 0 ≤ *p* ≤ 1. Changing the initial prey values only changes dynamics for the first few generations. Drawings above graphs illustrate the end behavior of the trajectories, where a red predator alone indicates SH fixes, a blue predator alone represents CH fixing, and a pair of blue and red predators together represent a polymorphic equilibrium. The parameters *β*_*s*_, depletion of small prey, *β*_*b*_, depletion of big prey, *b*_*s*_, benefit of small prey, *b*_*c*_, benefit of CH, and *H*, probability of horizontal transmission, written as (*β*_*s*_, *β*_*b*_, *b*_*s*_, *b*_*c*_, *H*), are **(a)** (0.2, 0.1, 0.1, 0.201, 0), **(b)** (0.2, 0.1, 0.1, 0.25, 0), **(c)** (0.9, 0.6, 0.15, 0.5, 0), **(d)** (0.5, 0.4, 0.09, 0.2, 0.1), **(e)** (0.5, 0.2, 0.2, 0.8, 0.25), **(f)** (0.5, 0.4, 0.2, 0.45, 0.5), **(g)** (0.5, 0.2, 0.2, 0.8, 0.1), and **(h)** (0.5, 0.3, 0.3, 0.8, 0.25).

If 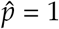 is stable, there is one polymorphic equilibrium, and we expect it to be unstable. From Result 3.2, this occurs if *β*_*b*_ < *γ*_1_ where *γ*_1_ can also be written as 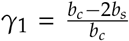. Thus there is one polymorphic equilibrium, and CH can fix for a wider range of BP depletion values if the BP is very beneficial and much more beneficial than SP.

If there are two polymorphic equilibria (i.e.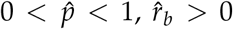, and 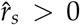) then 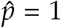 is unstable, and because 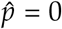 is stable (Result 3.3), we expect the larger polymorphic equilibrium to be stable and the smaller to be unstable. To confirm these predictions and to explore the stability of the polymorphic equilibrium, we compute the Jacobian *J*\* for local stability of the equilibrium 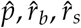, shown in (A2.6) - (A2.7). We created a parameter grid and evaluated the eigenvalues of the Jacobian for each parameter combination. As predicted, when there is one polymorphic equilibrium, it is unstable, and when there are two, the larger is stable. Thus the smaller polymorphic equilibrium determines the range of initial frequencies of CH *p*(0) for which CH will persist in the population (Fig. 3). The range of frequencies *p* of CH for which CH will persist is larger if ecological conditions favor CH: consuming BP is much more beneficial than small prey, i.e. 2*b*_*s*_/*b*_*c*_ is small, and depletion of SP *β*_*s*_ is high. However, low depletion of BP *β*_*b*_ only slightly increases this range.(Fig 3).

**Figure 3:**
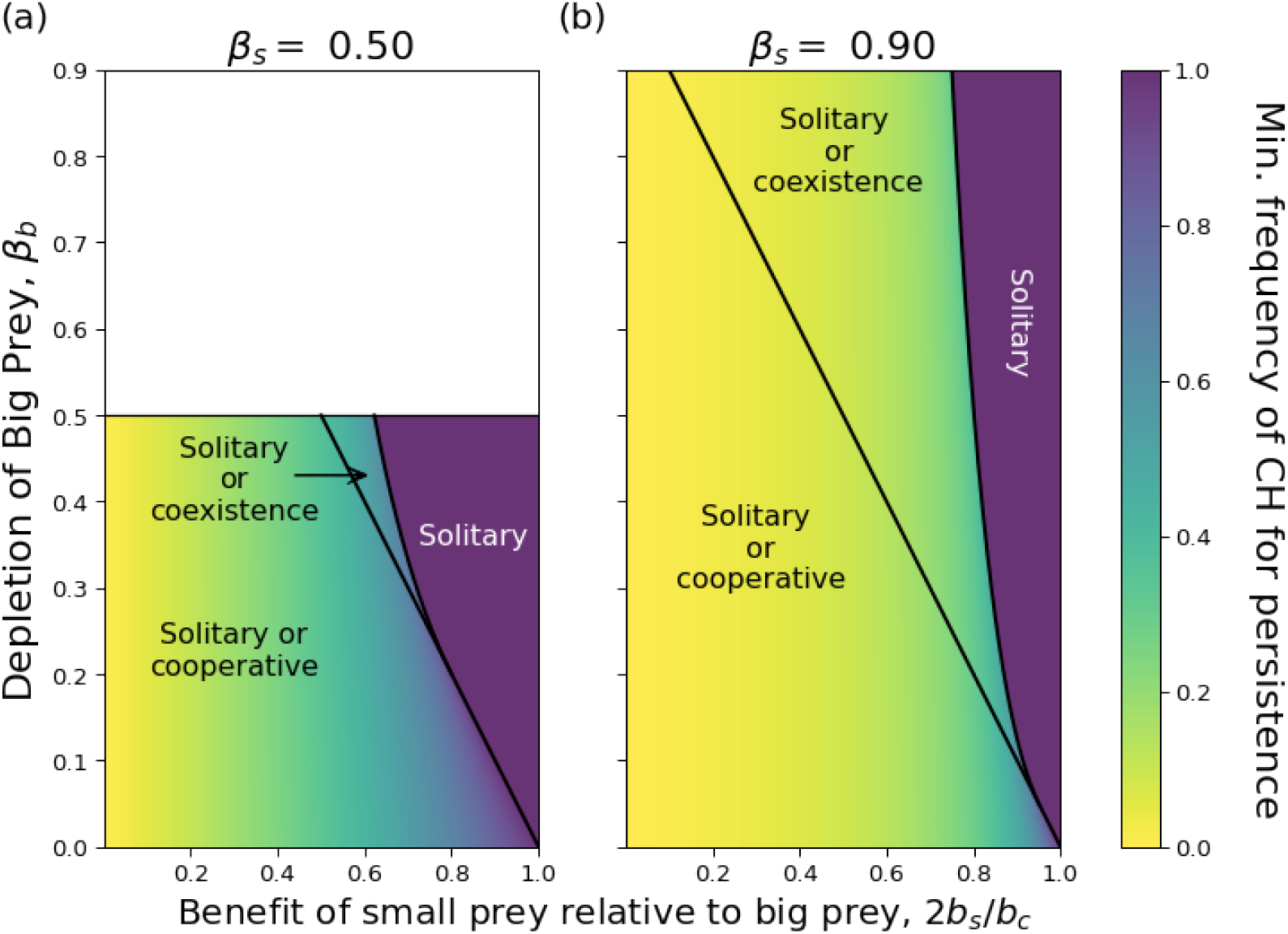
Without horizontal learning, *H* = 0, the minimum frequency of cooperative hunting (CH) such that CH persists (i.e. the minimum polymorphic equilibrium 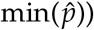) decreases if depletion of small, *β*_*s*_, is high and the benefit of small relative to BP, 2*b*_*s*_/*b*_*c*_, is low. The minimum equilibrium 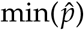 is shown for the small prey depletion constants (a) *β*_*s*_ = 0.5, and (b) *β*_*s*_ = 0.9. In the region labeled “Solitary or cooperative”, there is only one polymorphic equilibrium, which is unstable. In the region labeled “Solitary or coexistence” there are two polymorphic equilibria, and the one with the smaller 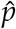 value is unstable. In the region labeled “Solitary”, for any starting frequency of CH, *p* < 1, CH will disappear from the population, i.e.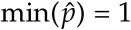.

### 3.2 Evolution of Cooperative Hunting with Horizontal Social Learning

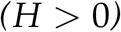

There can be zero, one, or two polymorphic equilibria 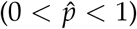, depending on the parameter values (Figs. 2d - 2f, A5.4, A5.5, A5.6). The conditions for existence of each of these equilibrium configurations are derived in Appendix (A3.1). The conditions for the existence of one and only one polymorphic equilibrium are: (a) horizontal learning *H* is high, the rate of depletion of BP *β*_*b*_ is low, availability 1 − *β*_*s*_ of SP is low, and the benefit of SP relative to BP 2*b*_*s*_/*b*_*c*_ is low, or (b) if the opposite conditions are true (i.e. *H* is low and *β*_*b*_, 1 − *β*_*s*_, and2*b*_*s*_/*b*_*c*_ are high). The conditions for the existence of either two or no polymorphic equilibria are more difficult to interpret.

Analysing the stability of the equilibria, we have the following results on the evolution of CH.

#### Result 3.5.

*CH increases when rare, i*.*e* 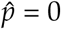 *is unstable, if (a) the BP is present, i*.*e*. 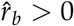, *but the SP is extinct, i*.*e*. 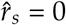, *or if (b) the prey species are at the equilibrium* 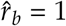 *and* 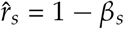, *and*

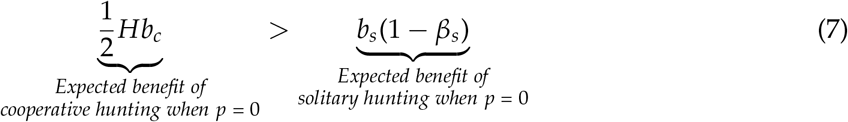

*Proof*. If *p* is close to 0, then from (2a), 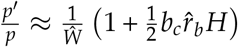, where from (3) 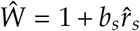 at 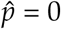
Then *p*′/*p* > 1 if 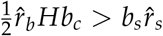. Thus, if 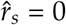 but 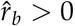, then CH increases when rare. If 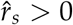, then since *p* ≈ 0 we have 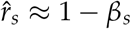 and 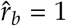. Then CH increases if 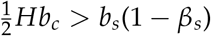.

The left side of (7) is the expected benefit of being a rare cooperative hunter because it is the benefit of big prey after sharing, 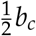, scaled by the probability of interacting with another cooperative hunter due to horizontal transmission, *H*. The right side is the expected benefit of SH when CH is rare because it is the benefit of small prey *b*_*s*_ scaled by the probability of obtaining small prey, 1 − *β*_*b*_. When CH is rare, the depletion of the BP due to CH is negligible, so that depletion of BP *β*_*b*_ does not affect the expected benefit of cooperative hunting.

Result 3.5 means that high enough probabilities of horizontal learning allow CH to evolve, even if hunting the SP is very beneficial (2*b*_*s*_/*b*_*c*_ close to 1) and the SP has a low depletion rate (*β*_*s*_ low).

#### Result 3.6.

*CH fixes (i*.*e*. 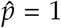 *is locally stable) if the BP is present, i*.*e*. 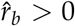, *and either (a) the SP is extinct, i*.*e*. 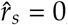, *or (b)* 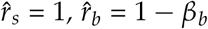 *and*

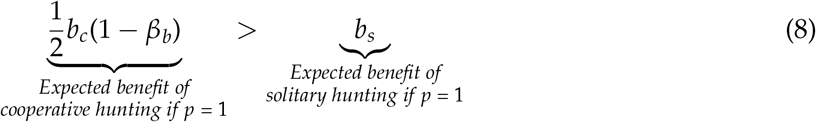

*Proof*. If *p* is very close to 1; i.e. *q* is very close to 0, then from (2b), 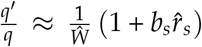 where from (3),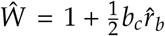. Then *q*′/*q* < 1 (and fixation of cooperative hunting is locally stable) if 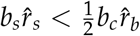. If the SP are extinct, i.e. *r*_*s*_ = 0, this inequality is trivially true. If SP are present, i.e. 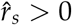, then 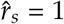. Also, 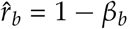, so *q*′/*q* < 1 if 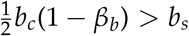.

Result 3.6 means that cooperative hunting cannot be invaded by solitary hunting if the BP has a low depletion rate (*β*_*b*_ low) and hunting the SP is much less beneficial than hunting the BP (2*b*_*s*_/*b*_*c*_ is small). Result 3.6 can also be interpreted as stating that solitary hunting invades if its fitness benefit is greater than the benefit of cooperative hunting when almost all predators are cooperators. Here, because solitary hunting is rare, the depletion constant of the SP, *β*_*s*_, does not affect the benefit of solitary hunting.

Note that inequality (8) is equivalent to *β*_*b*_ < *γ*_1_, the condition for CH to fix and one polymorphic equilibrium to exist if *H* = 0 (Result 3.2). CH fixes under the same conditions for *H* = 0 and *H* > 0 because horizontal transmission *H* does not enter in inequality (8).

#### 3.2.1 *Local Stability, H* > 0

For the local stability of a polymorphic equilibrium, we analyze the eigenvalues of the Jacobian *J*\* near the equilibrium 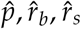, shown in (A3.4).

If there is no cycling, i.e. the eigenvalues are real, then we can use results 3.5 and 3.6, along with the conditions for zero, one or two polymorphic equilibria to exist (Appendix A3.1) to predict the local stability of polymorphic equilibria and suggest the following stability configurations:

1. There are no polymorphic equilibria and solitary hunting 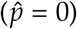 is the only stable equilibrium.
2. There are no polymorphic equilibria and cooperative hunting 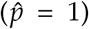 is the only stable equilibrium.
3. Neither solitary nor cooperative hunting fix (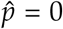 and 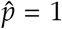 are not locally stable). There is one polymorphic equilibrium, which is stable.
4. Both solitary hunting and cooperative hunting can fix. There is one polymorphic equilibrium, which is unstable.
5. Solitary hunting can fix but cooperative hunting cannot. There are two polymorphic equilibria, the larger of which is stable.

Note that if SH cannot fix (CH invades) but CH can fix, there cannot be any polymorphic equilibria (see Appendix A3.1 for justification). Numerical calculations of the eigenvalues of *J*\* for 1000 randomly chosen parameter combinations confirmed that all equilibria fit with the five possibilities suggested above. The numerical calculations also confirmed results 3.5 and 3.6, i.e. that (a) CH increases when rare if the benefit of hunting the BP when CH is rare is greater than the benefit of hunting the SP when SH is common, and (b) that CH fixes if its benefit when common is greater than the benefit of SH when SH is rare.

#### 3.2.2 Convergent Stable Strategy of Cooperative Hunting

The convergent stable strategy, or CSS, of cooperative hunting is the smallest locally stable equilibrium frequency of CH. It is the stable strategy that can be reached from the cumulative evolution of small changes in the degree of cooperative hunting, as defined in Wakano and Aoki (2006). For example, if both 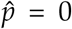 and 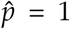 are locally stable, then the CSS is 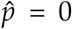. The probability of horizontal transmission *H*, the benefit ratio of SH to CH, 2*b*_*s*_/*b*_*c*_, and depletion of the SP, *β*_*s*_, all have large effects on the CSS, but the depletion rate of the BP, *β*_*b*_, only affects the CSS if the horizontal learning probability *H*, the benefit ratio 2*b*_*s*_/*b*_*c*_, and the depletion rate of SP *β*_*s*_ are large enough (Figs 4, A5.3).

**Figure 4:**
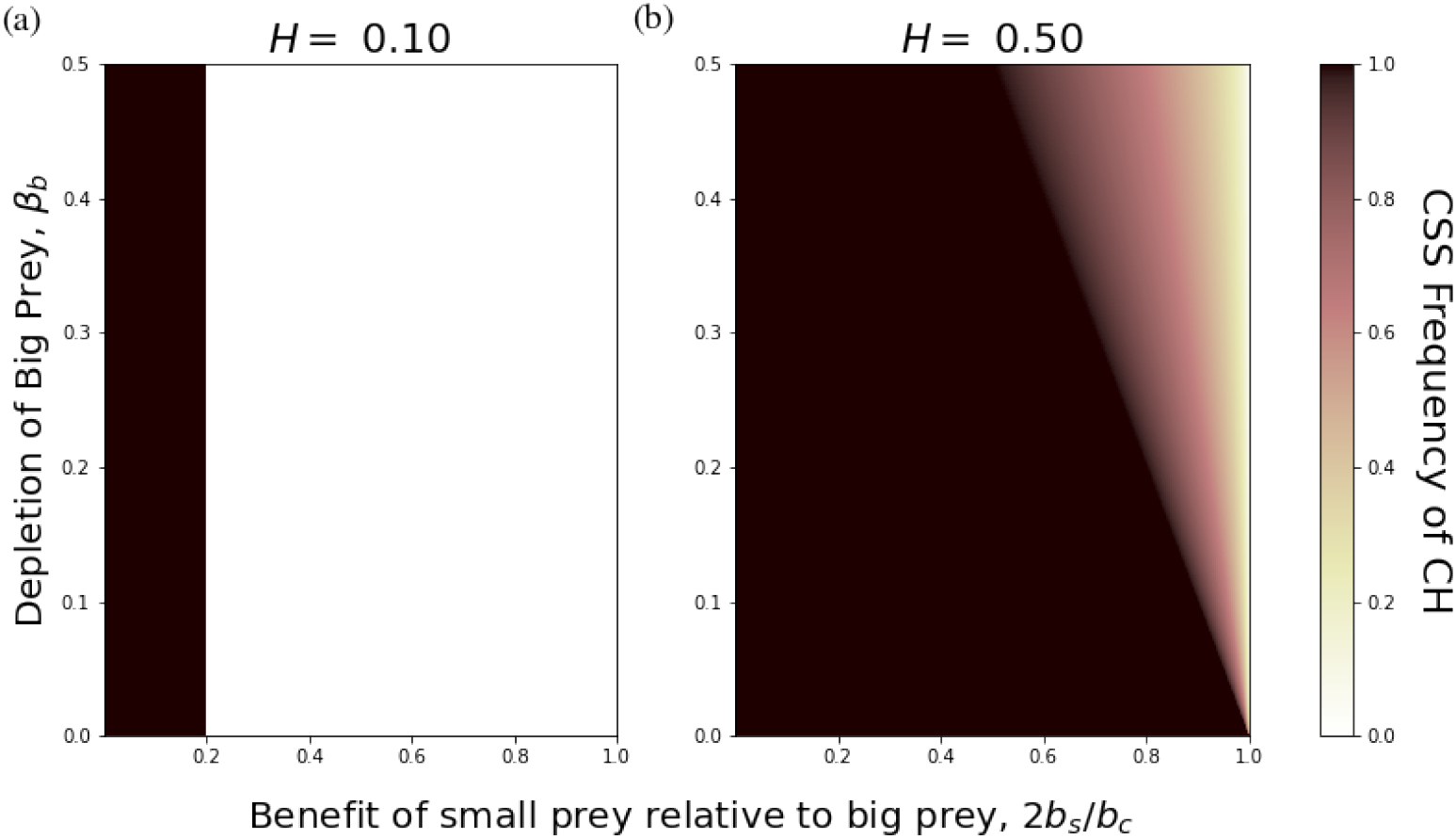
The convergent stable strategy, or CSS, of the frequency of cooperative hunting (shown by the color) versus the benefit ratio of solitary hunting to cooperative hunting, 2*b*_*s*_/*b*_*c*_ (x-axis) and the depletion of the big prey, *β*_*b*_ (y-axis), for depletion of small prey *β*_*s*_ = 0.5 and horizontal transmission probabilities **A)** *H* = 0.1 and **B)** *H* = 0.5. Cooperative hunting fixes in the black region and is lost in the white region.

At the CSS of the system, improving conditions for cooperative hunting, i.e. increasing horizontal learning *H*, decreasing the benefit ratio 2*b*_*s*_/*b*_*c*_, and lowering the depletion of big prey, *β*_*b*_, also increase population mean fitness, *W* (Fig A5.7). Comparing Fig. A5.7 to Fig. 4, population mean fitness of predators increases with the CSS frequency of cooperative hunting.

### 3.3 Evolution of Horizontal Social Learning, H

For the evolution of horizontal learning, we introduce a modifier locus that controls the probability of horizontal learning as in (Cohen et al., 2021; Feldman, 1972; Ram et al., 2018). This horizontal learning locus has alleles σ_1_ and σ_2_, which have probabilities of horizontal transmission *H* and *H* + *δ*_*H*_, respectively, with *δ*_*H*_ small and either positive or negative.

The two hunting behavior types are *ϕ* = *CH, SH*, cooperative and solitary hunting, respectively. The frequencies of σ_1_ and σ_2_ are *u* and *x*, respectively, and σ_1_ and σ_2_ are vertically transmitted independently of CH and SH. Let the frequencies of the four phenogenotypes in the offspring *CH*σ_1_, *SH*σ_1_, *CH*σ_2_, and *SH*σ_2_ be *u*_*c*_, *u*_*s*_, *x*_*c*_, *x*_*s*_, respectively, where *p* = *u*_*c*_ + *x*_*c*_ is the frequency of CH predators.

Adults interact and learn, with probabilities of adopting either CH or SH, for each interaction type shown in Table 2. The frequency of adults of type *CH*σ_1_, *SH*σ_1_, *CH*σ_2_, and *SH*σ_2_ after horizontal transmission are

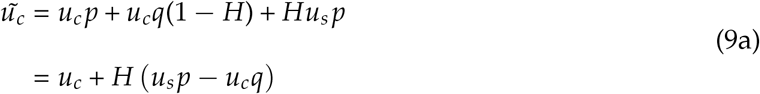

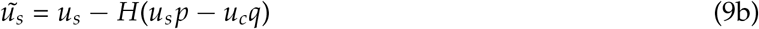

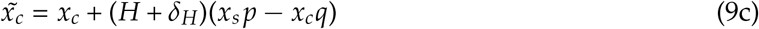

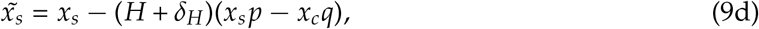

respectively. Interestingly, the quantity Δ = (*u*_*s*_ *p* − *u*_*c*_*q*) resembles a two-phenotype disequilibrium and could be regarded as a measure of covariance between the cooperative/solitary trait and the extent of learning (Ihara and Feldman, 2004). Note that 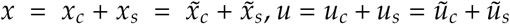,*u* = *u*_*c*_ + *u*_*s*_ = *ũ*_*c*_ + *ũ*_*s*_, but

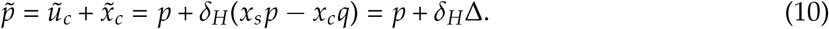

**Table 1:** Names and descriptions of the variables and parameters in the model

| Name | Definition |
| --- | --- |
| $p$ | frequency of cooperative hunting |
| $q$ | $1 - p$, the frequency of solitary hunting |
| $r_b$ | normalized density of the big prey |
| $r_s$ | normalized density of the small prey |
| $b_c$ | fitness benefit to the cooperative pair for hunting the big prey |
| $b_s$ | fitness benefit to an individual for hunting the small prey |
| $\beta_b$ | depletion constant by cooperative hunting of big prey |
| $\beta_s$ | depletion constant by solitary hunting of small prey |
| $H$ | the probability of horizontal learning from a potential hunting partner |

**Table 2:** Interactions between the four phenogenotypes if the horizontal learning mutant is present.

| Learner Type | Behavior of teacher | Frequency of encounter | Probability learner adopts hunting behavior |  |
| --- | --- | --- | --- | --- |
|  |  |  | CH | SH |
| $CH\sigma_1$ | CH | $u_cp$ | 1 | 0 |
| | SH | $u_cq$ | $1 - H$ | $H$ |
| $SH\sigma_1$ | CH | $u_sp$ | $H$ | $1 - H$ |
| | SH | $u_sq$ | 0 | 1 |
| $CH\sigma_2$ | CH | $x_cp$ | 1 | 0 |
| | SH | $x_cq$ | $1 - H - \delta_H$ | $H + \delta_H$ |
| $SH\sigma_2$ | CH | $x_sp$ | $H + \delta_H$ | $1 - H - \delta_H$ |
| | SH | $x_sq$ | 0 | 1 |

If *x* is very small (*x*_*s*_ and *x*_*c*_ are also very small), then increasing horizontal learning increases the frequency of cooperative hunting if *u*_*s*_ > *u*_*c*_, i.e. there are more solitary hunters than cooperators.

The fitness values of adults following these interactions are shown in Table 3. Then the frequency with which *CH*σ_1_ adults become parents is 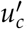, where

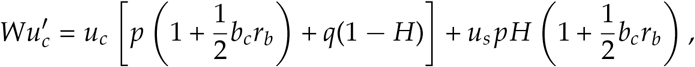

which simplifies to

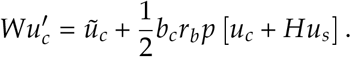

**Table 3:** Fitnesses of adults following hunting interactions.

| Learner Type | Behavior of teacher | Frequency of encounter | Expected fitness of behavior |  |
| --- | --- | --- | --- | --- |
|  |  |  | CH | SH |
| $CH\sigma_1$ | CH | $u_cp$ | $1 + \frac{1}{2}b_cr_b$ | 0 |
| | SH | $u_cq$ | $1 - H$ | $H(1 + b_sr_s)$ |
| $S\sigma_1$ | CH | $u_sp$ | $H(1 + \frac{1}{2}b_cr_b)$ | $(1 - H)(1 + b_sr_s)$ |
| | SH | $u_sq$ | 0 | $1 + b_sr_s$ |
| $CH\sigma_2$ | CH | $x_cp$ | $1 + \frac{1}{2}b_cr_b$ | 0 |
| | SH | $x_cq$ | $1 - H - \delta_H$ | $(H + \delta_H)(1 + b_sr_s)$ |
| $SH\sigma_2$ | CH | $x_sp$ | $(H + \delta_H)(1 + \frac{1}{2}b_cr_b)$ | $(1 - H - \delta_H)(1 + b_sr_s)$ |
| | SH | $x_sq$ | 0 | $1 + b_sr_s$ |

Thus the frequencies of *CH*σ_1_, *SH*σ_1_, *CH*σ_2_, and *SH*σ_2_ adults that become parents are given by the following recursions:

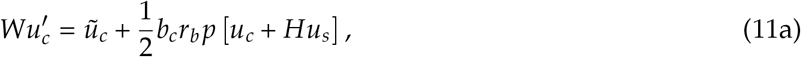

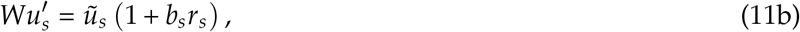

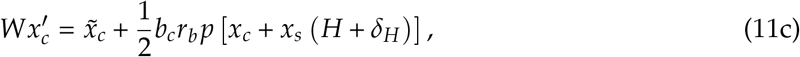

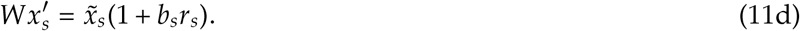

The population mean fitness is the sum of the right sides of (11), i.e.

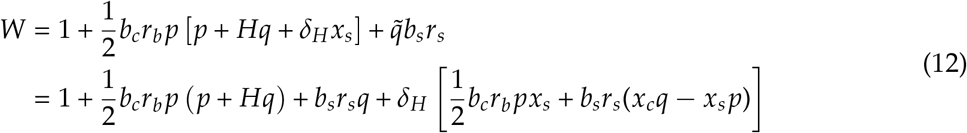

Assume the population is fixed on σ_1_. We assume the mutant σ_2_ appears near a locally stable polymorphic equilibrium 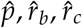 with 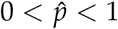 that solves (A3.2) (see Result 3.4 and Appendix A3.1 for parameter values that allow this equilibrium to exist), with 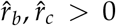. We disregard evolution of *H* near fixed solitary or cooperative hunting equilibria, 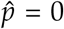 or 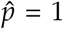 respectively, because if a hunting behavior is fixed in the population, horizontal learning cannot alter the learner’s behavior and thus will not affect evolution. Note that 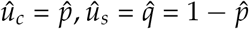.

#### Result 3.7.

*Phenotype* σ_2_, *with increased horizontal learning probability H* + *δ*_*H*_, *i*.*e. δ*_*H*_ > 0, *invades a population at the polymorphic equilibrium* 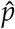, *which is fixed on* σ_1_ *with learning probability H and* 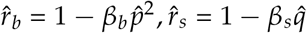.

*Proof*. The Jacobian *J*_*x*_ for linear increase of *x*_*c*_ and *x*_*s*_, whose eigenvalues determine whether σ_2_ invades, is shown in Appendix Eq. A4.2 and its eigenvalues are in Eq. A4.6. In Appendix A4, we show that the leading eigenvalue of this Jacobian is always greater than one.

Thus increased horizontal learning always invades a population in which both cooperative and solitary hunters are present, regardless of prey reward and depletion. It is important to note that when increased horizontal learning evolves, it also drives the evolution of increased CH (Fig 4) and can even lead to the polymorphic equilibria disappearing, causing CH to fix in the population (Figs A5.4, A5.5, A5.6). However, *CH* must be present in the system for *H* to evolve (because horizontal transmission does not affect cultural evolution if the population is fixed on SH, i.e., solitary hunting), but without horizontal transmission, *CH* must initially be present to reach a stable polymorphism.

## 4 Discussion

Horizontal learning allows cooperative hunting to evolve. Even if the benefit of small prey (i.e. *b*_*s*_) is small and competition for it is fierce and the big prey is plentiful and nutritious, CH cannot increase when initially rare without horizontal transmission (Result 3.2). This conclusion supports hypothesis (H1) but not hypotheses (H2) and (H3). Rare cooperative hunters cannot find a partner with which to hunt the big prey, and thus will not succeed unless solitary hunters can learn to cooperate. Similarly, in Cohen et al. (2021), high rates of horizontal transmission allowed a cooperative trait (in a Prisoner’s Dilemma game) to evolve. Horizontal learning helps a cooperator find a hunting partner (in our model) or increase the inclusive fitness of the cooperative phenotype (in Cohen et al. (2021)).

The result that cooperative hunting requires horizontal learning to evolve seems to contradict a game theory model (Packer and Ruttan, 1988), which concluded that cooperative hunting can evolve if the big prey is much more rewarding (e.g. more nutritious or very large and fatty) than the small prey. However, Packer and Ruttan (1988) analyzed the evolutionary stable strategies, the ESS, but did not discuss which ESS was actually attainable, i.e. whether cooperative hunting could increase when rare to the ESS frequency. Our model emphasizes the importance of identifying which evolutionary strategies are attainable and shows that an apparently optimal strategy may not actually emerge. Furthermore, we incorporate population dynamics of the prey, since depletion of prey can affect which hunting strategies have higher fitness.

Hypotheses (H2) and (H3) predict that strong competition for SP and weak competition for big prey should favor cooperative hunting. These hypotheses are supported if cooperative hunting is already present in the population, i.e. *p* is not very small, or if predators learn horizontally, *H* > 0. In these cases, if big prey are abundant and rewarding (*β*_*b*_ is low and *b*_*c*_ is high) compared to small prey (*β*_*s*_ is high and *b*_*s*_ is low), then cooperative hunting can spread through the population and fix, resulting in all predators becoming cooperators. However, from analysis of the CSS of the system, the relative importance of each of these cultural and environmental factors in decreasing order would be: probability of horizontal transmission > benefit ratio of the prey types, 2*b*_*s*_/*b*_*c*_ and depletion rate of small prey, *β*_*s*_ > depletion rate of big prey, *β*_*b*_. The impact of prey abundance on predator cooperation may be seen in wolves: while the grey wolves of North America hunt in large packs, Arabian wolves in the Middle East have been observed to hunt in groups of only 2 - 4 individuals. The much smaller pack size has been attributed to the extinction of large prey in the Middle East and abundance of human trash (which does not require cooperation) that wolves can scavenge (Hefner and Geffen, 1999).

Our results predict that a species that can hunt both solitarily and cooperatively should be able to learn not only from parents, but also from peers within the same generation. Such social learning may not require the ability to copy, but may occur through social facilitation (Giraldeau and Caraco, 2018) if predators are socially attentive enough to notice a conspecific hunting, are willing to join a hunt, and can share the catch. Further research on animals such as the fossas discussed earlier (Lührs and Dammhahn, 2010) may clarify the directionality of cultural transmission, i.e. whether it occurs horizontally, vertically, or obliquely from non-parental adults.

Introducing a mutant with an increased tendency to learn horizontally produced two surprising results. First, the horizontal learning and hunting strategy traits are not independent during the stage where juveniles interact and can learn hunting behaviors from each other. Horizontal learning acts to control the force of phenotypic disequilibrium, analogous to the rate of recombination in population genetics models (Ihara and Feldman, 2004). However, unlike the prediction of hypothesis (H4), at the polymorphic equilibrium, increased horizontal learning evolves irrespective of any of the other model parameters. Horizontal learning allows a predator to match its behavior to that of its interaction partner, which benefits the cooperator and does not harm a solitary hunter. This result may change if horizontal learning of the solitary behavior were to result in solitary hunters directly interfering with each other, e.g. if the learner tried to steal the prey that an unwitting demonstrator was hunting.

Here, horizontal learning can only evolve from stable polymorphism (i.e. both cooperative and solitary hunting are present in the population), but the population may be fixed at one of these hunting behaviors. The evolution of horizontal learning may be significant, even if all individuals use the same hunting strategy, if predators not only learn foraging strategies, but also prey locations, horizontally. For example, ravens likely share information about the location of food at communal roosts (Wright et al., 2003) and honeybees use the waggle dance to recruit each other to particular food sites (I’Anson Price and Grüter, 2015). In our model, a predator that learns horizontally to hunt solitarily is assumed to watch a current solitary hunter and then proceed to hunt a small prey by itself. However, if horizontal learning were to communicate the location of food, then a producer-scrounger game might emerge between solitary hunters. In a producer-scrounger game, some predators, the producers, individually discover food while others, the scroungers, watch the producers and steal some of the food the producers find (Giraldeau and Caraco, 2018). Cooperative hunting may be able to evolve even if the benefit of sharing the big prey is less than the benefit of solitarily catching the small prey, i.e.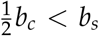, if solitary hunters that learn from each other were to compete directly for found food.

Predictions from our research may be valuable for conservation and wildlife reintroduction programs. For example, a program to reintroduce the Asiatic wild ass (*Equus hemionus*) in Israel began in 1983 (Saltz and Rubenstein, 1995), and in recent years this population has become established. Our model suggests that the new availability of a large ungulate, for example this ass, may enable Arabian wolves to hunt more cooperatively, which could translate to larger pack sizes.

Our model analyzes predator behavior frequencies, but not predator population density. To further understand the ecological interactions underpinning the evolution of cooperative hunting, future studies should incorporate the population dynamics of the predator. Although cooperative hunting and horizontal learning can result in a high frequency of predators performing the same behavior, in our model neither of the prey types went extinct; the 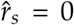 and 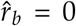 equilibria were each unstable. This may not be the case if predator population densities change. If the increase in fitness from cooperative hunting causes a dramatic increase in predator population density, then the more valuable big prey may go extinct. These analyses can guide wildlife reintroduction programs including that of the aforementioned Asiatic wild ass.

Social and cooperative predators such as killer whales, wolves, some species of sea otters (Eisenberg, 2013; Estes et al., 1998), and humans play an essential role in structuring ecosystems across the globe. This study advances a nascent field (e.g. see (Berec, 2010; Borofsky and Feldman, 2022) that studies the influence of social learning and cooperation by predators on ecosystems, and emphasizes that there is a feedback loop between prey availability and the degree of cooperation among predators.

## 5 Acknowledgements

We thank Eli Geffen, Amiyaal Ilany, Ofek Kraus, Arnon Lotem, Ran Barkai, and Tal Simon for discussions and comments. This work was supported in part by the Council for Higher Education of Israel’s PhD Sandwich Scholarship Program (TMB), Morrison Institute for Population and Resource Studies at Stanford University (MWF), John Templeton Foundation (MWF and YR), Israel Science Foundation (YR 552/19), and the Minverva Shiftung Center for Lab Evolution (YR).

## Notes

### Competing Interest Statement

The authors have declared no competing interest.

### Summary of Updates

Fixing typos and clarifying sentence structure; adding dotted lines to appendix figures A5.2, A5.4 - A5.6; adding additional trajectory panels to figure 2 to show all stability configurations; explaining that cannot have polymorphic equilibria if cooperation invades and fixes; deleting the omega variable in appendix figures and captions because it is not necessary; removing unnecessary and confusing colorbar in appendix figure A5.3; honeybee example in discussion; reference to Ihara and Feldman (2004)

