## Appendices for "Cultural transmission, competition for prey, and the evolution of cooperative hunting"

#### A1 Prey Densities

The prey densities are $N_s$ and $N_b$, respectively, and their carrying capacities are $K_s$ and $K_b$, respectively. For simplicity we assume $K_s = K_b = K$. Thus, the normalized densities of the prey types are $r_s = N_s/K$ and $r_b = N_b/K$, respectively.

The number of SP in the next generation is

$$N_s' = \frac{N_s(e_s - f_s(1-p))}{c_s + d_s N_s} \tag{A1.1}$$

where $e_s$, $c_s$, and $d_s$ are non-negative constants describing SP's population size-dependent growth. The predation term is $f_s(1-p)$, where $f_s > 0$ is the strength of solitary predation (encompassing the rate of encountering single predators) and $1-p$ is the probability that a single predator is a solitary hunter. We can change variables from population size to a normalized density between 0 and 1, as in Borofsky and Feldman (2022): the carrying capacity of SP is $K = (e_s - c_s)/d_s$. Then the normalized density (here on simply referred to as density) of small prey is $r_s = N_s/K = (N_s d_s)/(e_s - c_s)$, and (A1.1) becomes

$$r_s' = \frac{r_s\left[1 + \eta_s - \beta_s(1-p)\right]}{1 + r_s\eta_s}\ , \tag{A1.2}$$

where $\eta_s = e_s/c_s - 1 > 0$ is SP's growth constant, and $\beta_s = f_s/c_s$ is SP's predation depletion constant. For simplicity, we assume $\eta_s = 1$.

The density of BP in the next generation is

$$N_b' = \frac{N_b(e_b - f_b p^2)}{c_b + d_b N_b}\ , \tag{A1.3}$$

where $e_b$, $c_b$, and $d_b$ are non-negative constants that determine the BP's population size-dependent growth. The predation term is $f_b p^2$ where $f_b$ is the encounter rate with a pair of predators and $p^2$ is the probability that both of these predators are cooperative. We assume $f_b < f_s$. The carrying capacity of big prey is $K = (e_b - c_b)/d_b$ and it is assumed that $e_b/c_b > 1$ and $d_b(e_s - c_s) = d_s(e_b - c_b)$). Then $r_b = (N_b d_b)/(e_b - c_b)$ and (A1.3) becomes

$$r'_b = \frac{r_b\left[1 + \eta_b - \beta_b p^2\right]}{1 + r_b \eta_b} , \tag{A1.4}$$

where $\eta_b = e_b/c_b - 1 > 0$ is BP's growth constant, and $\beta_b = f_b/c_b$, is BP's predation depletion constant. We assume $\beta_b < \beta_s < 1$ because BP need to encounter two predators at once to be caught and for simplicity set $\eta_b = 1$ (i.e. $e_b/c_b = 2$) and thus $\eta_b = \eta_s = 1$.

### A2 Vertical Transmission Only, $H = 0$

#### *A2.1 Proof of Result 3.2*

To find the polymorphic equilibrium, divide Eqs. 2a by 2b, substituting $H = 0$, i.e.,

$$1 = \frac{1 + \hat{p}\frac{b_c \hat{r}_b}{2}}{1 + b_s \hat{r}_s} ,$$

giving

$$b_s \hat{r}_s = \frac{\hat{p} b_c \hat{r}_b}{2} . \tag{A2.1}$$

Next, substitute $\hat{r}_s = 1 - \beta_s(1 - \hat{p})$ and $\hat{r}_b = 1 - \beta_b \hat{p}^2$, into Eq. A2.1 to give the cubic equation

$$0 = -\beta_b \hat{p}^3 + \hat{p}\left(1 - \beta_s \frac{2b_s}{b_c}\right) - \frac{2b_s}{b_c}(1 - \beta_s) = Q(p) . \tag{A2.2}$$

Because $Q(0) < 0$ and as $p \to -\infty$, $Q(p) \to +\infty$, $Q(p)$ must have at least one root that is negative. At $p = 1$,

$$Q(1) = 1 - \beta_b - \frac{2b_s}{b_c} .$$

If $Q(1) > 0$, i.e. $b_c(1-\beta_b) > 2b_s$, then the cubic has one root between 0 and 1. This completes the first part of the proof, namely that there is one polymorphic equilibrium if $\beta_b < \gamma_1$. If $Q(1) < 0$, then the $Q(\cdot)$ could have 0 or two roots between 0 and 1. However, the critical points of $Q(p)$ are at

$$p^*_\pm = \pm\sqrt{\frac{1-\frac{2b_s}{b_c}\beta_s}{3\beta_b}} = \pm\sqrt{M},$$

where $M = \frac{1-\frac{2b_s}{b_c}\beta_s}{3\beta_b}$. If $Q(1) > 0$ (i.e. $b_c(1-\beta_b) > 2b_s$), $0 < p^*_+ \leq 1$ and $Q(p^*_+) < 0$, then there are two polymorphic equilibria.

Note that $p^*_+ > 0$ is real by our assumption that $b_c > 2b_s$. Then $p^*_+ < 1$ if

$$3\beta_b > 1 - \frac{2b_s}{b_c}\beta_s, \tag{A2.3}$$

or equivalently if $\beta_b > \gamma_2$. Furthermore,

$$\begin{aligned} Q(p^*_+) &= \sqrt{M}\left(-\beta_b M + 1 - b\beta_s\right) - \frac{2b_s}{b_c}(1-\beta_s) \\ &= \frac{2}{3}\sqrt{M}\left(1 - \frac{2b_s}{b_c}\beta_s\right) - \frac{2b_s}{b_c}(1-\beta_s)\,, \end{aligned} \tag{A2.4}$$

and since $b_c > 2b_s\beta_s$, $Q(p^*_+) > 0$ if

$$\sqrt{M} > \frac{3b_s(1-\beta_s)}{b_c(1-2\frac{b_s}{b_c}\beta_s)},$$

i.e.,

$$\left(1 - \frac{2b_s}{b_c}\beta_s\right)^3 > 27\beta_b\left(\frac{b_s}{b_c}\right)^2(1-\beta_s)^2, \tag{A2.5}$$

or equivalently, $\beta_b < \gamma_3$.

#### A2.2 Stability of Polymorphic Equilibrium

The Jacobian $J^*$ for local stability of any equilibrium $\hat{p}, \hat{r}_b, \hat{r}_s$ is of the form

$$J^* = \begin{pmatrix} \frac{\partial p'}{\partial p} & \frac{\partial p'}{\partial r_b} & \frac{\partial p'}{\partial r_s} \\ \frac{\partial r_b'}{\partial p} & \frac{\partial r_b'}{\partial r_b} & 0 \\ \frac{\partial r_s'}{\partial p} & 0 & \frac{\partial r_s'}{\partial r_s} \end{pmatrix} \left((p, r_b, r_s) = (\hat{p}, \hat{r}_b, \hat{r}_s)\right) , \tag{A2.6}$$

where

$$\frac{\partial p'}{\partial p} = \frac{1 + \hat{p}(b_s \hat{r}_s + b_c \hat{r}_b (1 - \hat{p}))}{\hat{W}} \tag{A2.7a}$$

$$\frac{\partial p'}{\partial r_b} = \frac{\frac{1}{2} b_c \hat{p}^2 (1 - \hat{p})}{\hat{W}} \tag{A2.7b}$$

$$\frac{\partial p'}{\partial r_s} = -\frac{\hat{p}}{\hat{W}} b_s (1 - \hat{p}) \tag{A2.7c}$$

$$\frac{\partial r_b'}{\partial p} = -2 \frac{\beta_b \hat{p} \hat{r}_b}{1 + \hat{r}_b} \tag{A2.7d}$$

$$\frac{\partial r_b'}{\partial r_b} = \frac{2 - \beta_b \hat{p}^2}{(1 + \hat{r}_b)^2} \tag{A2.7e}$$

$$\frac{\partial r_s'}{\partial p} = \frac{\beta_s \hat{r}_s}{1 + \hat{r}_s} \tag{A2.7f}$$

$$\frac{\partial r_s'}{\partial r_s} = \frac{2 - \beta_s (1 - \hat{p})}{(1 + \hat{r}_s)^2} \ . \tag{A2.7g}$$

Note that if $0 < \hat{p} < 1, \hat{r}_b = 1 - \beta_b \hat{p}^2$, $\hat{r}_s = 1 - \beta_s (1 - \hat{p})$, and $\hat{W} = 1 + \frac{1}{2} b_c \hat{p} \hat{r}_b = 1 + b_s \hat{r}_s$. Then

$$\frac{\partial p'}{\partial r_b} = \frac{\hat{p}^2 b_c (1 - \hat{p})}{2\hat{W}} \tag{A2.8}$$

$$\frac{\partial r_b'}{\partial r_b} = \frac{1}{1 + \hat{r}_b} \tag{A2.9}$$

$$\frac{\partial r_s'}{\partial r_s} = \frac{1}{1 + \hat{r}_s}, \tag{A2.10}$$

where the partial derivatives are evaluated at the equilibrium ($\hat{p}, \hat{r}_b = 1 - \beta_b \hat{p}^2, \hat{r}_s = 1 - \beta_s(1 - \hat{p})$).

### A3 Horizontal Learning $H > 0$

#### *A3.1 Equilibria with $H > 0$*

To find the polymorphic equilibrium $\hat{p}$, divide 2a by 2b, giving

$$b_s \hat{r}_s = \frac{1}{2} b_c \hat{r}_b (\hat{p} + H(1 - \hat{p})). \tag{A3.1}$$

Substituting $\hat{r}_s = 1 - \beta_s(1 - \hat{p})$ and $\hat{r}_b = 1 - \beta_b \hat{p}^2$ in (A3.1), the equilibria are roots of the cubic equation

$$0 = p^3 \beta_b (H - 1) - H\beta_b p^2 + p\left(1 - H - \frac{2b_s}{b_c}\beta_s\right) + H - \frac{2b_s}{b_c}(1 - \beta_s). \tag{A3.2}$$

Call the cubic on the right side of (A3.2) $Q_H(p)$. The leading coefficient $\beta_b(H-1) < 0$ so as $p \to \infty$, $Q_H(p) \to -\infty$. At $p = 0$,

$$Q_H(0) = H - \frac{2b_s}{b_c}(1 - \beta_s),$$

which is positive if $H > \frac{2b_s}{b_c}(1 - \beta_s)$. At $p = 1$,

$$Q_H(1) = 1 - \beta_b - \frac{2b_s}{b_c}.$$

The critical points of $Q_H$ are

$$p^*_{+,-} = \frac{H\beta_b \pm \sqrt{\beta_b \left\{H^2\beta_b + 3(1 - H)\left[1 - H - \beta_s(2b_s/b_c)\right]\right\}}}{3\beta_b(H - 1)}. \tag{A3.3}$$

For parameter values such that $p^*$ is real, then $p^*_+ < 0$ because the denominator $\beta_b(H-1)$ is negative, and $p^*_- > 0$ for parameter values such that $1 - H - \beta_s(2b_s/b_c) < 0$. Thus we have the following types of equilibria:

1. Only one valid polymorphic equilibrium exists if either
   (a) $Q_H(0) < 0$ and $Q_H(1) > 0$, i.e. horizontal learning $H$ is high, the rate of depletion of the big prey $\beta_b$ is low, availability $1-\beta_s$ of the small prey is low, and the benefit of the small prey relative to the big prey $2b_s/b_c$ is low. From Results 3.5 and 3.6, this means that there is one polymorphic equilibrium if cooperation can increase when rare and cannot fix if it becomes common.
   (b) $Q_H(0) > 0$ and $Q_H(1) < 0$, which are the opposite ecological conditions to (a). From Results 3.5 and 3.6, this means that there is one polymorphic equilibrium if cooperation cannot increase when rare (i.e. if $p$ starts below the polymorphic equilibrium, then it goes to $\hat{p} = 0$) and can fix if common (i.e. if $p$ starts above the polymorphic equilibrium, then it goes to $\hat{p} = 1$).

   Note that existence of three real roots of (A3.2) is not possible because if $p^*_{+,-}$ from (A3.3) are real, then at least one of the critical points is negative.

2. There are two polymorphic equilibria if
   (a) $Q_H(0), Q_H(1) < 0$, i.e. cooperation cannot invade or fix. Note that if $Q_H(0), Q_H(1) > 0$ (i.e. cooperation invades and fixes) then there cannot be two polymorphic equilibria because the leading coefficient of $Q_H(p)$ is negative and the left critical point is $p^*_+ < 0$.
   (b) The rightmost critical point of $Q_H$, called $p^*_-$ is between 0 and 1.
   (c) $Q_H(p^*_-) < 0$.

3. Otherwise, if sign($Q_H(0)$) = sign($Q_H(1)$) and sign($Q_H(p^*_-)$ = sign($Q_H(0)$) or $p^*_-$ is not both real and between 0 and 1, then there are no polymorphic equilibria. This

means that for any starting frequency of cooperation, either cooperation disappears ($p$ goes to $\hat{p} = 0$), or it fixes ($p$ goes to $\hat{p} = 1$).

#### *A3.2 Local Stability with $H > 0$*

We use linear stability analysis to evaluate stability of the $(\hat{p}, \hat{q}, \hat{r}_s, \hat{r}_b)$ equilibria to perturbations in the frequency of CH and generate Figures A5.3 A5.4, A5.5, A5.6, A5.7. This Jacobian for local stability of the equilibrium $\hat{p}$ (from solving (A3.2)), $\hat{r}_b$ , $\hat{r}_s$ is of the form

$$J^* = \left.\begin{pmatrix} \frac{\partial p'}{\partial p} & \frac{\partial p'}{\partial r_b} & \frac{\partial p'}{\partial r_s} \\ \frac{\partial r_b'}{\partial p} & \frac{\partial r_b'}{\partial r_b} & 0 \\ \frac{\partial r_s'}{\partial p} & 0 & \frac{\partial r_s'}{\partial r_s} \end{pmatrix}\right|_{(\hat{p},\hat{r}_b,\hat{r}_s)} . \tag{A3.4}$$

From (2), $p'$ is a function of $r_b, p$ and $W$, which is in turn a function of $p, r_b, r_s$. Thus the partial derivatives of $p'$ depend on those of $W$. The partial derivatives of the population mean fitness $W$ with respect to $p$, $r_b$, and $r_s$ at the polymorphic equilibrium. These are

$$\frac{\partial W}{\partial p} = \frac{1}{2} b_c \hat{r}_b \left[2p(1-H) + H\right] - b_s \hat{r}_s \tag{A3.5a}$$

$$\frac{\partial W}{\partial r_b} = \frac{1}{2} \hat{p} b_c (\hat{p} + H\hat{q}) \tag{A3.5b}$$

$$\frac{\partial W}{\partial r_s} = \hat{q} b_s. \tag{A3.5c}$$

From (2a),

$$\frac{\partial p'}{\partial p} = \frac{1}{\hat{W}} \left\{ 1 + \frac{1}{2} b_c \hat{r}_b (1-\hat{p}) \left[2\hat{p}(1-H) + H\right] + \hat{p} b_s \hat{r}_s \right\} . \tag{A3.6}$$

Differentiating (2a) with respect to $r_b$,

$$\frac{\partial p'}{\partial r_b} = \frac{1}{2\hat{W}} b_c \hat{p}(1-\hat{p}) \left[\hat{p} + H(1-\hat{p})\right] \tag{A3.7}$$

and differentiating (2a) with respect to $r_s$

$$\frac{\partial p'}{\partial r_s} = -\frac{1}{\hat{W}^2}\frac{\partial W}{\partial r_s}\hat{W}\hat{p} = \frac{-\hat{p}\hat{q}b_s}{\hat{W}}. \tag{A3.8}$$

The partial derivatives $\frac{\partial r_b'}{\partial p}$, $\frac{\partial r_b'}{\partial r_b}$, $\frac{\partial r_s'}{\partial p}$, and $\frac{\partial r_s'}{\partial r_s'}$ are the same as in (A2.7 d - g).

### A4 Evolution of Horizontal Learning, $H$

For Jacobian $J_x$, we write $x_c'$, $x_s'$ for $x_c, x_s$ very small:

$$\begin{aligned} x_c' &= \frac{1}{\hat{W}}\left\{x_c + (H+\delta_H)(x_s\hat{p} - x_c\hat{q}) + \frac{1}{2}b_c\hat{r}_b\hat{p}\left[x_c + x_s(H+\delta_H)\right]\right\} \\ x_s' &= \frac{1}{\hat{W}}\left[x_s + (H+\delta_H)(x_c\hat{q} - x_s\hat{p})\right](1 + b_s\hat{r}_s)\,, \end{aligned}$$

which, when gathered in terms of $x_c$, $x_s$, are

$$\begin{aligned} x_c' &= x_c\frac{1}{\hat{W}}\left[1 + \frac{1}{2}b_c\hat{r}_b\hat{p} - q(H+\delta_H)\right] + x_s\frac{\hat{p}}{\hat{W}}(H+\delta_H)\left(1 + \frac{1}{2}b_c\hat{r}_b\right) \\ x_s' &= x_s\frac{1}{\hat{W}}\left[1 - p(H+\delta_H)\right](1 + b_s\hat{r}_s) + x_c\frac{\hat{q}}{\hat{W}}(H+\delta_H)(1 + b_s\hat{r}_s)\,. \end{aligned}$$

Thus the Jacobian $J_x$ is

$$J_x = \begin{pmatrix} \frac{1}{\hat{W}}\left[1 + \frac{1}{2}b_c\hat{r}_b\hat{p} - \hat{q}(H+\delta_H)\right] & \frac{\hat{p}}{\hat{W}}(H+\delta_H)\left(1 + \frac{1}{2}b_c\hat{r}_b\right) \\ \frac{\hat{q}}{\hat{W}}(H+\delta_H)(1 + b_s\hat{r}_s) & \frac{1}{\hat{W}}\left[1 - \hat{p}(H+\delta_H)\right](1 + b_s\hat{r}_s) \end{pmatrix}, \tag{A4.1}$$

where

$$\hat{W} = 1 + \hat{p}\frac{1}{2}b_c\hat{r}_b(\hat{p} + H\hat{q}) + \hat{q}b_s\hat{r}_s,$$

from (3). Note that if $\hat{p}, \hat{q} \neq 0$, then $\hat{W} = 1 + \frac{1}{2} b_c \hat{r}_b (\hat{p} + H\hat{q})$ and $\hat{W} = 1 + b_s \hat{r}_s$ from (2), so the Jacobian is

$$J_x = \begin{pmatrix} \frac{1}{\hat{W}} \left[ 1 + \frac{1}{2} b_c \hat{r}_b \hat{p} - \hat{q}(H + \delta_H) \right] & \frac{\hat{p}}{\hat{W}} (H + \delta_H) \left( 1 + \frac{1}{2} b_c \hat{r}_b \right) \\ \hat{q}(H + \delta_H) & [1 - \hat{p}(H + \delta_H)] \end{pmatrix}, \tag{A4.2}$$

which has trace

$$Tr(J_x) = 1 - \hat{p}(H + \delta_H) + \frac{1 + \frac{1}{2} b_c \hat{r}_b \hat{p} - \hat{q}(H + \delta_H)}{1 + b_s \hat{r}_s} \tag{A4.3}$$

and determinant

$$\begin{aligned} \det(J_x) &= \frac{1}{\hat{W}} \left[ 1 + \frac{1}{2} b_c \hat{r}_b \hat{p} - (H + \delta_H) \left( 1 + \frac{1}{2} b_c \hat{r}_b \hat{p}^2 \right) - \frac{1}{2} b_c \hat{r}_b \hat{p} \hat{q} \, (H + \delta_H)^2 \right] \\ &= \frac{1}{\hat{W}} \left\{ 1 - H + \frac{1}{2} b_c \hat{r}_b \hat{p} \left[ 1 - H \left( \hat{p} + \hat{q} H \right) \right] - \delta_H \left( 1 + \frac{1}{2} b_c \hat{r}_b \hat{p}^2 + 2 \hat{p} \hat{q} \right) - \mathcal{O}\left( \delta_H^2 \right) \right\}. \end{aligned} \tag{A4.4}$$

Note that for $\delta_H$ very small, the determinant must be positive.

The eigenvalues of $J_x$ are solutions to the characteristic polynomial

$$0 = \lambda_x^2 - Tr(J_x)\lambda_x + \det(J_x), \tag{A4.5}$$

and are

$$\lambda_x = \frac{1}{2} \left\{ Tr(J_x) \pm \sqrt{Tr(J_x)^2 - 4\det(J_x)} \right\}. \tag{A4.6}$$

At $\lambda = 1$, the characteristic polynomial is

$$\begin{aligned} Q_x(1) &= \hat{p} \left( H + \delta_H \right) + \frac{1}{\hat{W}} \left[ \left( q - 1 - \frac{1}{2} b_c \hat{r}_b \hat{p}^2 \right) \left( H + \delta_H \right) - \frac{1}{2} b_c \hat{r}_b \hat{p} \hat{q} \left( H + \delta_H \right)^2 \right] \\ &= \hat{p} \left( H + \delta_H \right) \left\{ 1 - \frac{1}{\hat{W}} \left[ 1 + \frac{1}{2} b_c \hat{r}_b \hat{p} + \frac{1}{2} b_c \hat{r}_b \hat{q} \left( H + \delta_H \right) \right] \right\}, \end{aligned}$$

but since $\hat{W} = 1 + \frac{1}{2}b_c\hat{r}_b\hat{p} + \frac{1}{2}b_c\hat{r}_b H\hat{q}$ from (2),

$$
\begin{aligned}
Q_x(1) &= \hat{p}\left(H + \delta_H\right)\left[1 - \frac{1}{\hat{W}}\left(\hat{W} + \frac{1}{2}b_c\hat{r}_b\hat{q}\delta_H\right)\right] \\
&= -\frac{1}{\hat{W}}\left(H + \delta_H\right)\frac{1}{2}b_c\hat{r}_b\hat{p}\hat{q}\delta_H
\end{aligned}
$$

which is negative if $\delta_H > 0$. Since $Q_x(\lambda)$ opens upward and at $\lambda = 0$, $Q_x(0) = \det(J_x) > 0$, there must be an eigenvalue greater than one. Thus any polymorphic equilibrium that is stable in the absence of $x_c$ and $x_s$ is unstable to invasion by a mutant with increased horizontal learning.

### A5 Figures

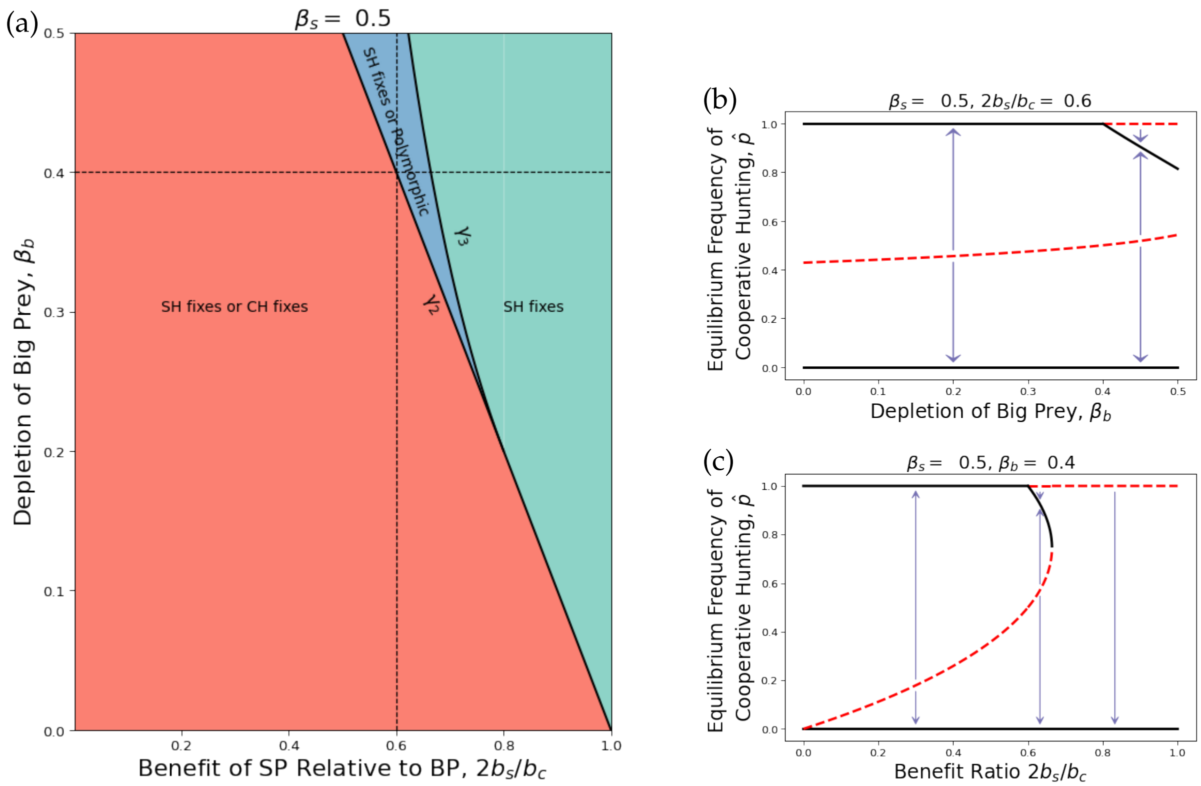


Figure A5.1: Evolution of cooperative hunting under vertical transmission only (i.e. $H = 0$) if the rate of depletion of the SP is intermediate ($\beta_s = 0.5$). **(a)** Ranges of the benefit ratio of hunting the SP versus the BP, $2b_s/b_c$ (x-axis) and depletion of the BP, $\beta_b$ (y-axis) for global fixation of solitary hunting (turquoise), bistability of fixation on solitary hunting and coexistence of cooperative hunting and solitary hunting at $p = \hat{p}_2$ (blue), and bistability of fixation on solitary and cooperative hunting (red). The vertical dotted line in **(a)** indicates the slice shown in the bifurcation diagram in panel **(b)** and the horizontal dotted line in **(a)** indicates the slice shown in the bifurcation diagram in panel **(c)**. In Panels **(b)** and **(c)**, black lines indicate stable equilibria of the frequency of cooperative hunting $\hat{p}$ and red dotted lines are unstable equilibria. In panel **(b)** the x-axis is $\beta_b$, for $\frac{2b_s}{b_c} = 0.6$, and in panel **(c)** the x-axis is $b$, for $\beta_b = 0.4$. We see that the magnitude of stable $\hat{p} > 0$ increases with lower $\beta_b$ and $b$.

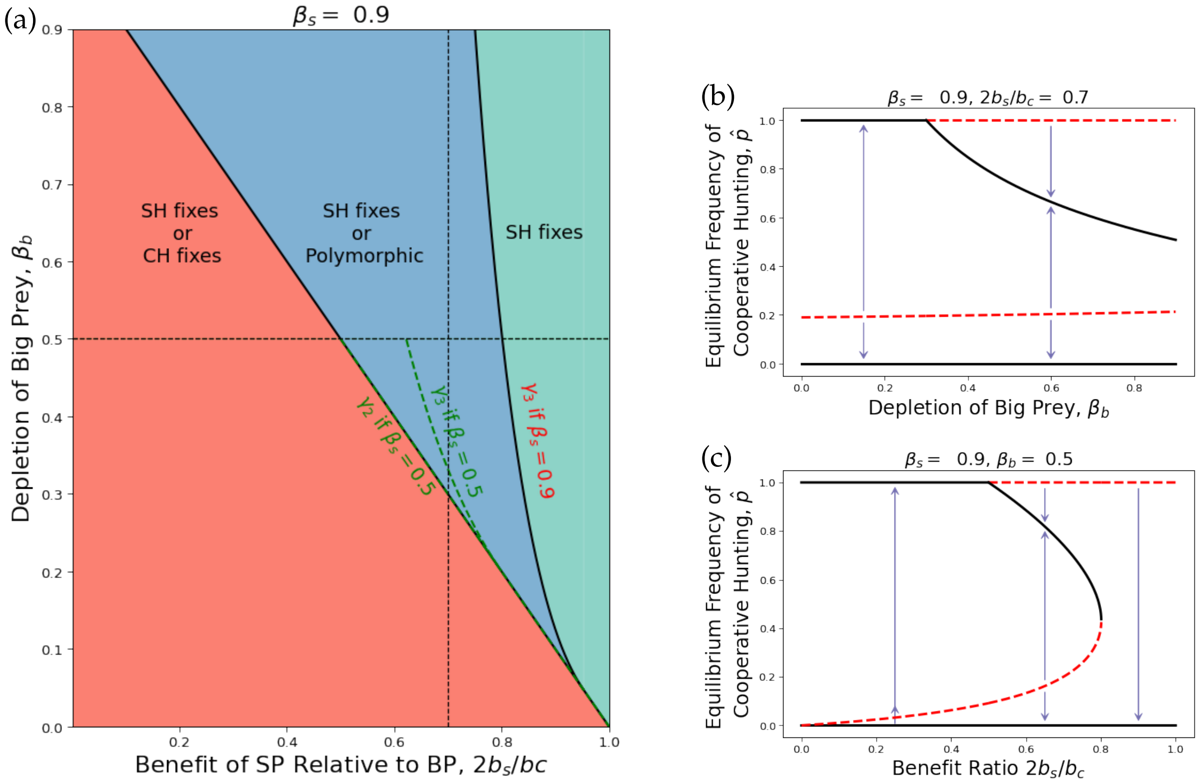


Figure A5.2: Evolution of cooperative hunting under vertical transmission only (i.e. $H = 0$) if the rate of depletion of the SP is high ($\beta_s = 0.9$). **(a)** Parameter ranges of the benefit ratio of hunting the SP versus the BP, $2b_s/b_c$ (x-axis) and depletion of the BP, $\beta_b$ (y-axis) for global fixation of solitary hunting (turquoise), bistability of fixation on solitary hunting and coexistence of cooperative hunting and solitary hunting at $p = \hat{p}_2$ (blue), and bistability of fixation on solitary and cooperative hunting (red). The vertical dotted line indicates the slice shown in the bifurcation diagram in panel **(b)** and the horizontal dotted line indicates the slice shown in the bifurcation diagram in panel **(c)**. In Panels **(b)** and **(c)**, black lines indicate stable equilibria of the frequency of cooperative hunting $\hat{p}$ and red dotted lines are unstable equilibria. In panel **(b)** the x-axis is $\beta_b$, for $\frac{2b_s}{b_c} = 0.7$, and in panel **(c)** the x-axis is $b$, for $\beta_b = 0.5$. We see that the magnitude of stable $\hat{p} > 0$ increases with lower $\beta_b$ and $b$.

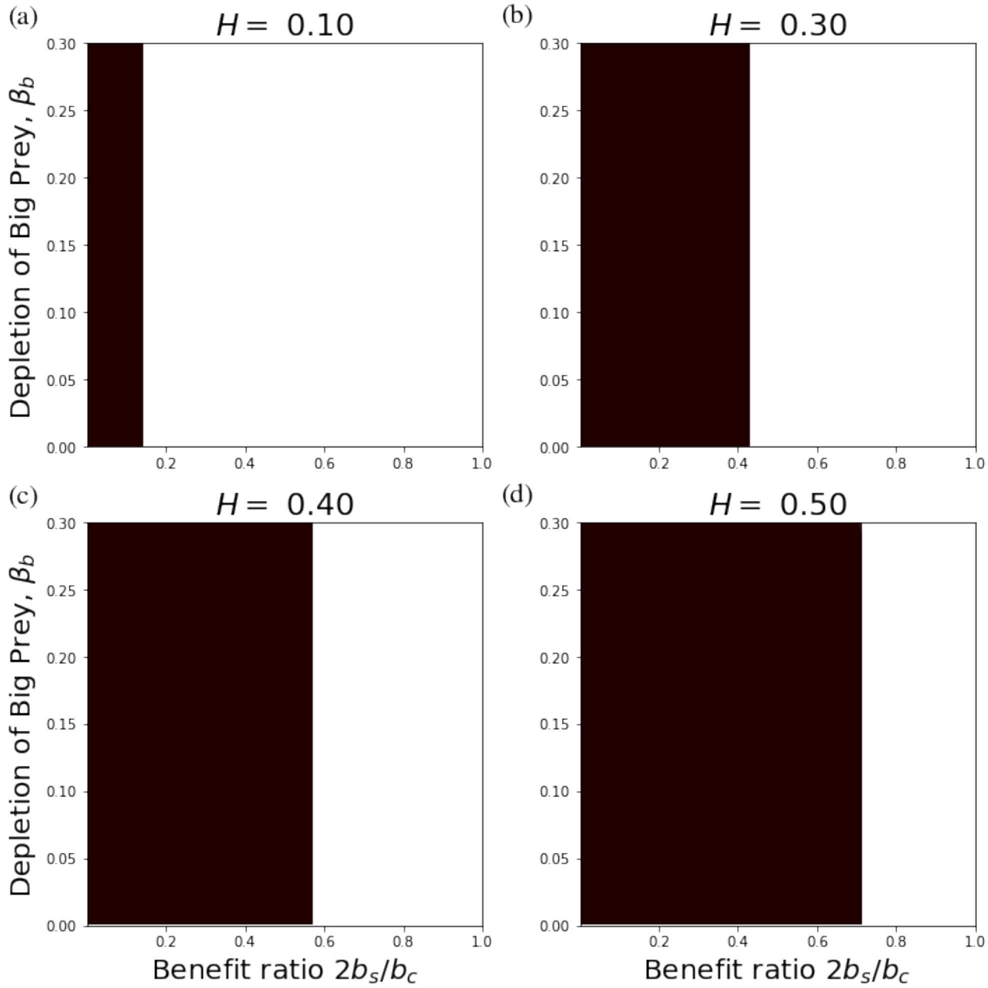


Figure A5.3: The convergent stable strategy (CSS) frequency of CH versus the benefit ratio of solitary hunting to cooperative hunting, $2b_s/b_c$ (x-axis) and the depletion of the big prey, $\beta_b$ (y-axis), for depletion of small prey $\beta_s = 0.3$ and horizontal transmission probabilities **A)** $H = 0.1$, **B)** $H = 0.3$, **C)** $H = 0.4$, and **D)** $H = 0.5$. Cooperative hunting fixes in the black region and is lost in the white region.

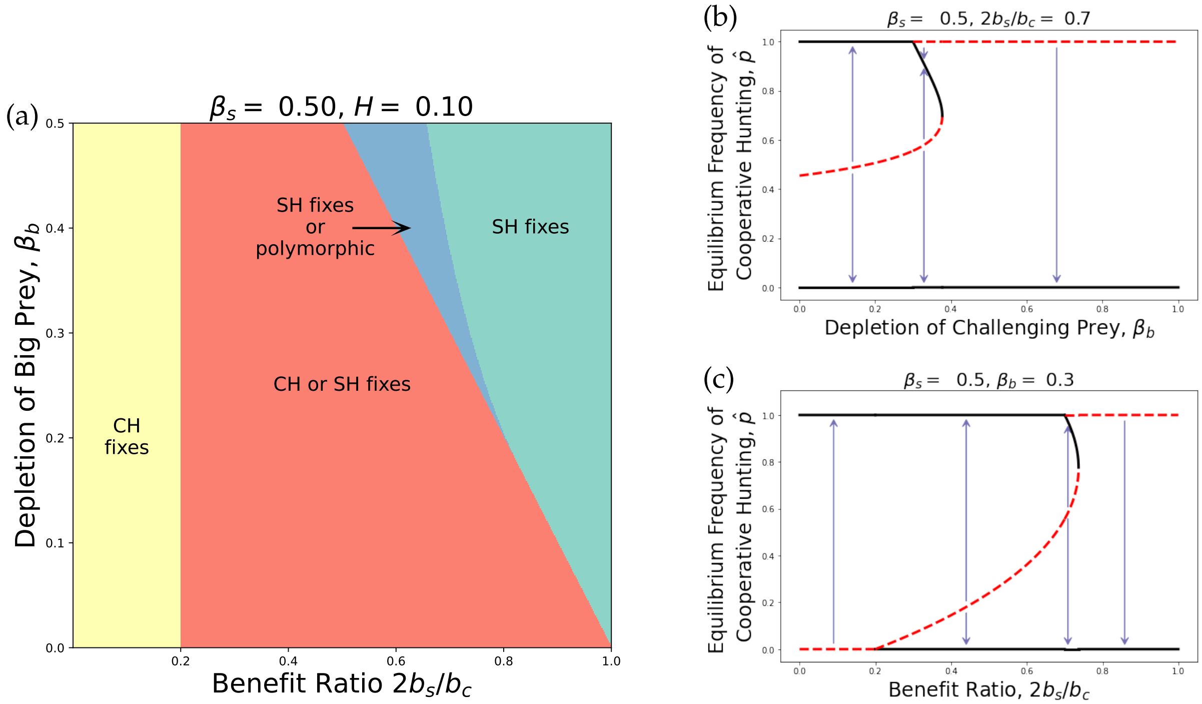


Figure A5.4: Evolution of cooperative hunting with horizontal learning ($H = 0.1$) if the rate of depletion of the SP is moderate ($\beta_s = 0.5$). **(a)** Parameter ranges of the benefit ratio of hunting the SP versus the BP, $2b_s/b_c$ (x-axis) and depletion of the BP, $\beta_b$ (y-axis) for global fixation of cooperative hunting (CH; yellow), bistability of fixation on solitary hunting (SH) or CH (red), bistability of fixation on CH or coexistence of CH and SH (blue), and global fixation on SH (turquoise). The vertical dotted line in **(a)** indicates the slice shown in the bifurcation diagram in panel **(b)** and the horizontal dotted line in **(a)** indicates the slice shown in the bifurcation diagram in panel **(c)**. In Panels **(b)** and **(c)**, black lines indicate stable equilibria of the frequency of cooperative hunting $\hat{p}$ and red dotted lines are unstable equilibria. In panel **(b)** the x-axis is $\beta_b$, for $b = 2b_s/b_c = 0.8$, and in panel **(c)** the x-axis is $2b_s/b_c$, for $\beta_b = 0.3$. We see that the magnitude of stable $\hat{p} > 0$ increases with lower $\beta_b$ and $2b_s/b_c$.

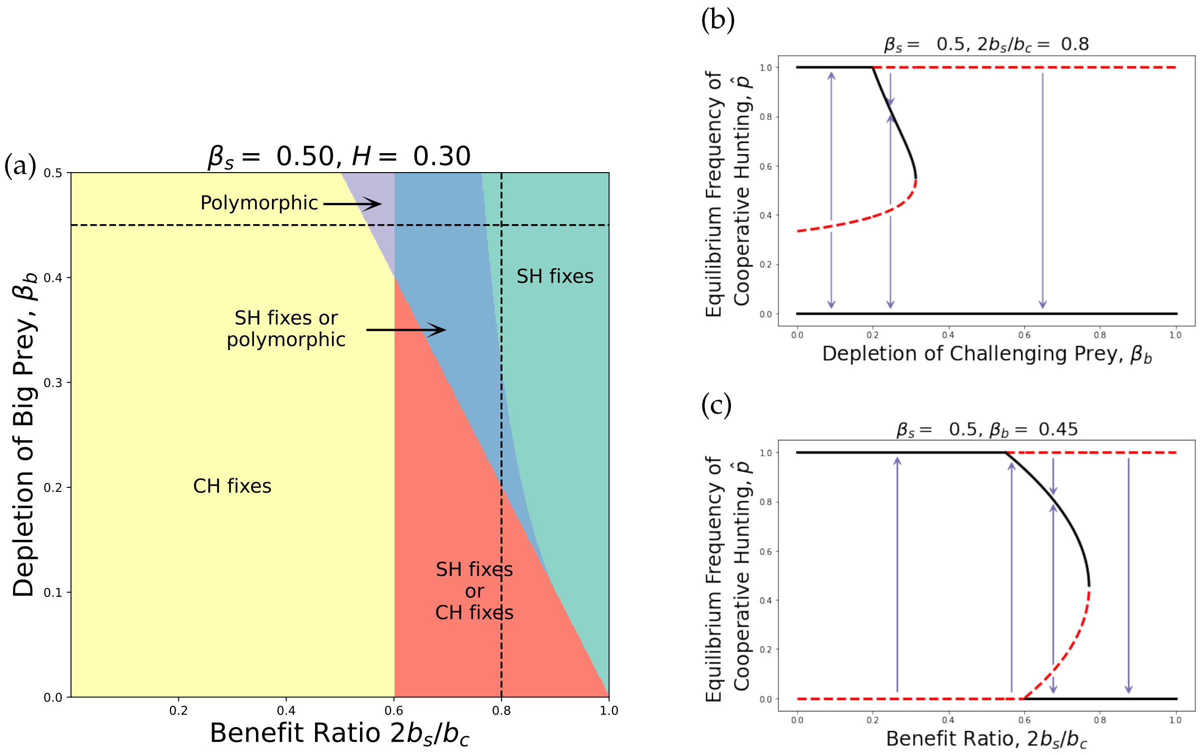


Figure A5.5: Evolution of cooperative hunting with horizontal learning ($H = 0.3$) if the rate of depletion of the SP is moderate ($\beta_s = 0.5$). **(a)** Parameter ranges of the benefit ratio of hunting the SP versus the BP, $2b_s/b_c$ (x-axis) and depletion of the BP, $\beta_b$ (y-axis) for global fixation of cooperative hunting (CH; yellow), bistability of fixation on solitary hunting (SH) or CH (red), bistability of fixation on CH or coexistence of CH and SH (blue), global fixation on SH (turquoise), and stability of coexistence of CH and SH (purple). The vertical dotted line in **(a)** indicates the slice shown in the bifurcation diagram in panel **(b)** and the horizontal dotted line in **(a)** indicates the slice shown in the bifurcation diagram in panel **(c)**. In Panels **(b)** and **(c)**, black lines indicate stable equilibria of the frequency of cooperative hunting $\hat{p}$ and red dotted lines are unstable equilibria. In panel **(b)** the x-axis is $\beta_b$, for $2b_s/b_c = 0.8$, and in panel **(c)** the x-axis is $2b_s/b_c$, for $\beta_b = 0.45$. We see that the magnitude of stable $\hat{p} > 0$ increases with lower $\beta_b$ and $2b_s/b_c$.

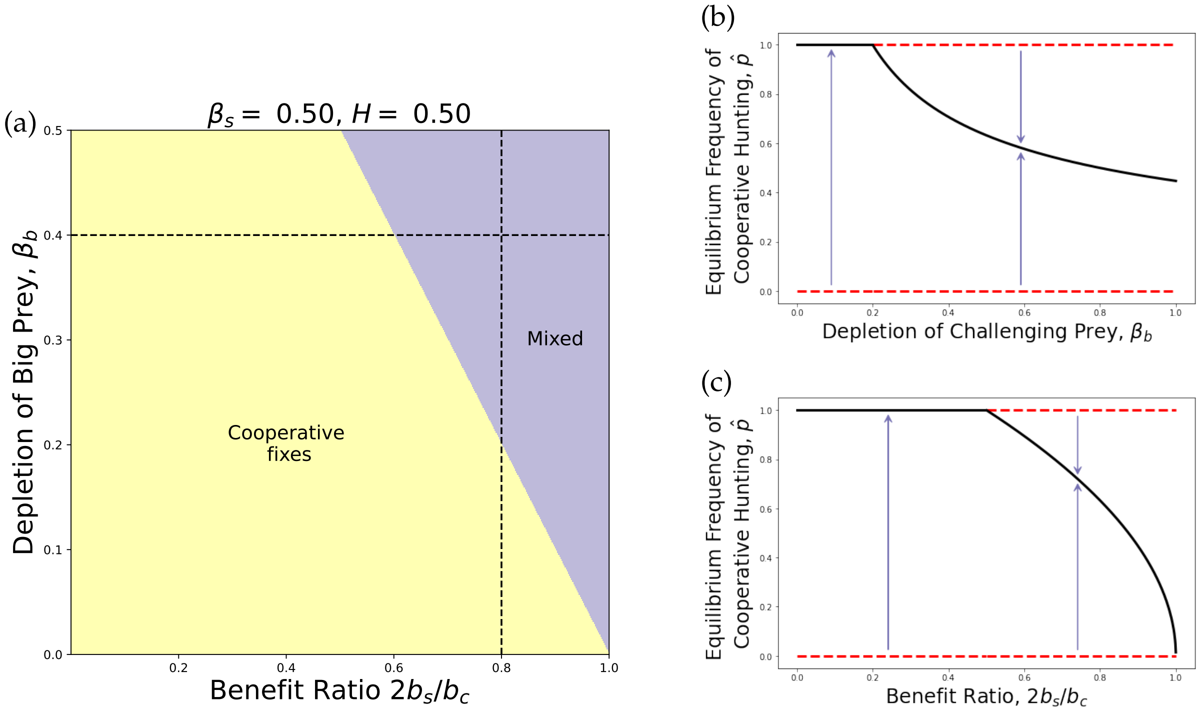


Figure A5.6: Evolution of cooperative hunting with horizontal learning ($H = 0.5$) if the rate of depletion of the SP is moderate ($\beta_s = 0.5$). **(a)** Parameter ranges of the benefit ratio of hunting the SP versus the BP, $2b_s/b_c$ (x-axis) and depletion of the BP, $\beta_b$ (y-axis) for global fixation of cooperative hunting (CH; yellow) and stability of coexistence of CH and SH (purple). The vertical dotted line in **(a)** indicates the slice shown in the bifurcation diagram in panel **(b)** and the horizontal dotted line in **(a)** indicates the slice shown in the bifurcation diagram in panel **(c)**. In Panels **(b)** and **(c)**, black lines indicate stable equilibria of the frequency of cooperative hunting $\hat{p}$ and red dotted lines are unstable equilibria. In panel **(b)** the x-axis is $\beta_b$, for $2b_s/b_c = 0.8$, and in panel **(c)** the x-axis is $2b_s/b_c$, for $\beta_b = 0.4$. We see that the magnitude of stable $\hat{p} > 0$ increases with lower $\beta_b$ and $2b_s/b_c$.

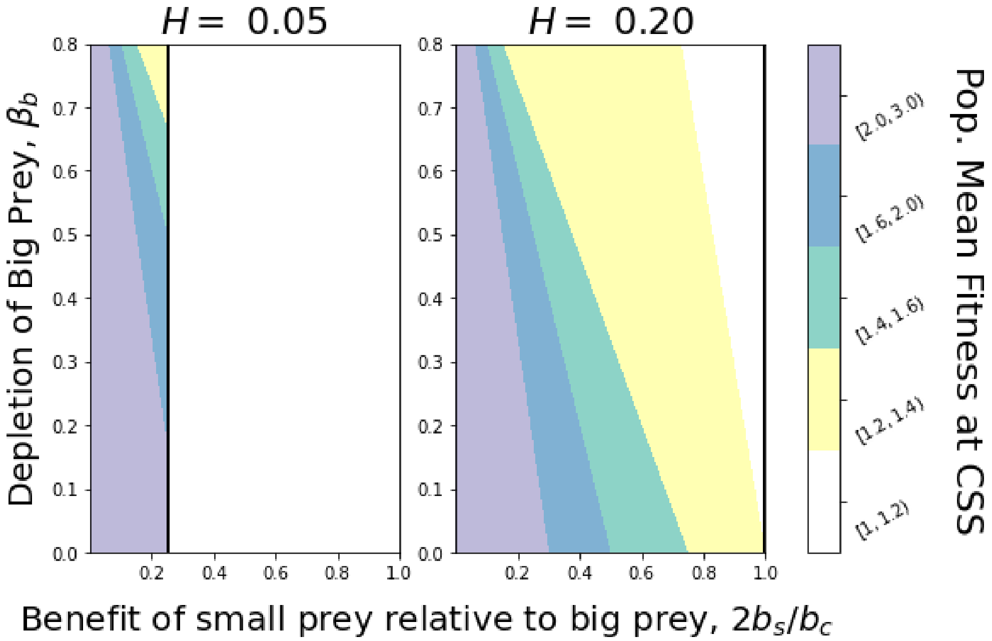


Figure A5.7: Population mean fitness, $\hat{W}$, at the convergent stable strategy, CSS, of cooperative hunting, $\hat{p}$, for $\beta_s = 0.8, b_s = 0.3$. Population mean fitness is colored by assigning the $\hat{W}$ at each parameter combination to one of the five bins ($[1, 1.2), [1.2, 1.4), [1.4, 1.6), [1.6, 2.0)$, and $[2.0, 3.0]$), as described in the colorbar. The vertical black line indicates the cutoff value of $2b_s/b_c$ for the invasion of cooperative hunting (see Result 3.5); to the left of this line, cooperative hunting can increase from rarity.
